# Neural signatures of human psychological resilience driven by acute stress

**DOI:** 10.1101/2024.03.24.586503

**Authors:** Noriya Watanabe, Shinichi Yoshida, Ruedeerat Keerativittayayut, Masaki Takeda

## Abstract

Neurophysiological mechanisms underlying psychological resilience, the ability to overcome adversity^1,2^, have been extensively studied in animals. However, in comparison with animals, human resilience is unique in that it is underpinned by higher cognitive functions, such as self-confidence and a positive attitude to challenges^3,4^. Given these discrepancies, the neurophysiological mechanisms underlying human resilience remain unclear. To address this issue, we recorded multimodal responses after acute stress exposure over 1.5 hours using functional brain imaging and peripheral physiological measurements. Here, we showed that the degree of individual resilience is indexed by multiple changes in neural dynamics 1 hour after acute stress. Both functional magnetic resonance imaging and electroencephalography show that activity in the cortical salience network and power in high-beta and gamma oscillations increase in less resilient individuals. Contrastingly, activity in the cortical default mode network and spontaneous activity in the posterior hippocampus increase in more resilient individuals. Machine learning analysis confirmed that, 1 hour after stress exposure, the functional connectivity in the salience network was the most influential, followed by that in the default mode network, gamma power, high-beta power, and hippocampal activity. The neurophysiological dynamics for resilience do not occur as previously thought, but rather in a time-lagged manner against stress exposure. Our findings Shed light on a new approach to recovery from stress-induced deficits such as delayed neuromodulation after a stressful event.

---

Psychological resilience is an individual ability to adapt successfully to acute stress, trauma, or chronic forms of adversity^1,2^. Despite its critical role in mental health, this ability varies considerably among individuals (Fig. 1a). Those with lower resilience are at high risk of depression and other mental illnesses^3,5^. Accumulating evidence has demonstrated that acute stress drives successive responses in the brain and body, such as rapid autonomic^6-15^, time-lagged (peaking 20 min after) stress-hormonal^16-24^, and immune and slow-corticosteroid responses (peaking more than 1 hour after) (Fig. 1b)^25-30^. These homeostatic responses may contribute to the initiation of the processes underlying resilience.

**Fig. 1:**
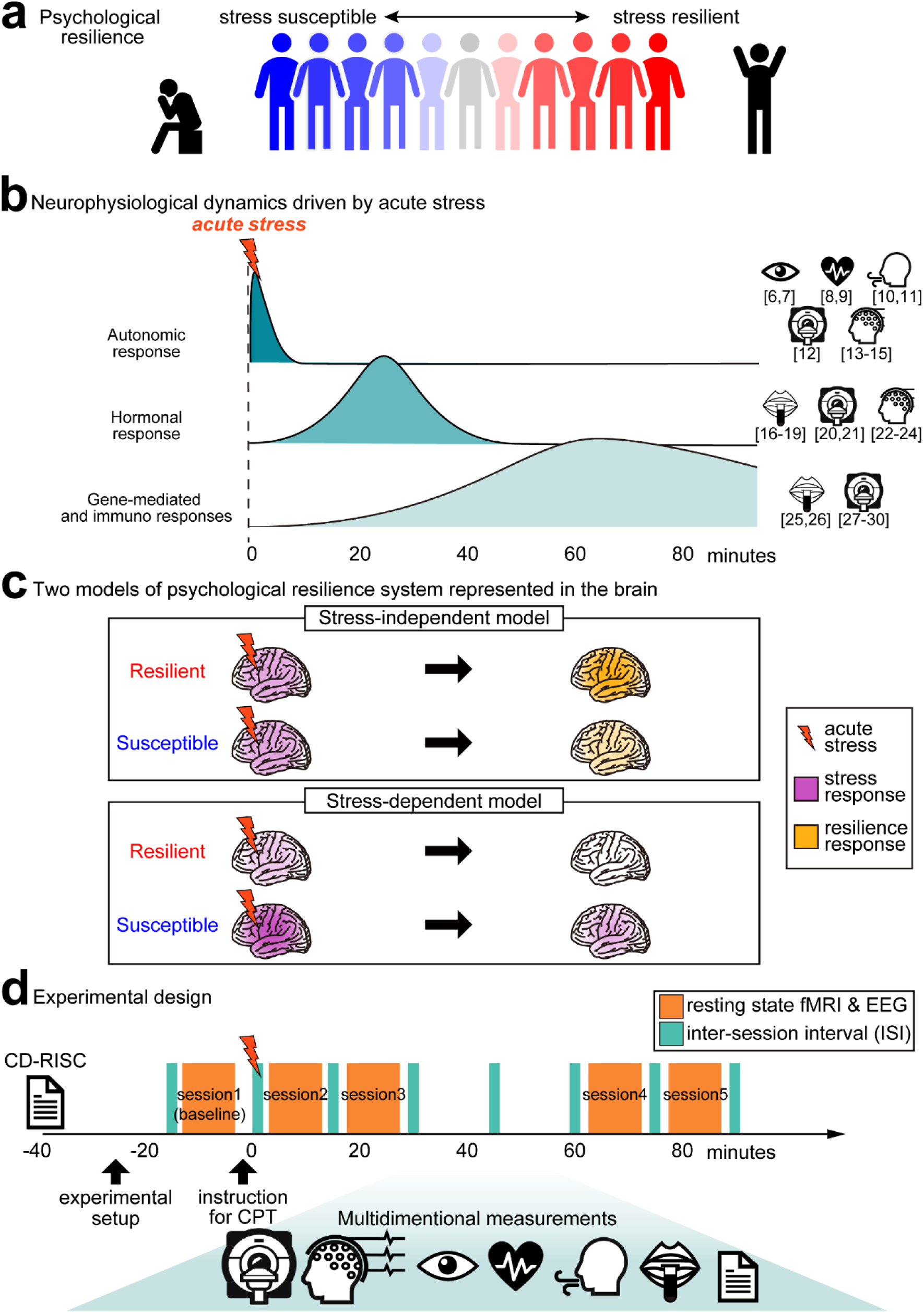
Hypothesis and experimental design. (**a**) Diversity of psychological resilience. Individuals with low and high adaptability to stressful environments are referred to as susceptible and resilient, respectively. (**b**) Three stress-related neurophysiological responses to acute stress from previous studies. The first response is a transient autonomic nervous system response during the stress experience, such as pupil dilation, increased heart and respiration rates, and changes in brain state as measured by fMRI and EEG. The second response is the stress hormone response, including the increase in salivary cortisol and changes in brain state, which peak 20 minutes after the stress event. The third response is the immune and slow-cortisol response, which lasts more than 1 hour and is thought to affect brain activity. (**c**) The two hypothesised models for resilience-related neurophysiological responses. The first model hypothesises unique resilience responses independent of sensitivity to the stressor; the second model, stress-dependent resilience responses. We aimed to identify the substantive model by assessing whether the responses would be correlated with individual resilience levels but not with the stress responses shown in (**b**) (the former model), or would be associated with both individual resilience and stress responses (the latter model). (**d)** The current experiment examined acute stress-related neurophysiological responses over 1 hour. First, each participant completed the resilience-related questionnaires. Then, six types of neurophysiological responses (fMRI, EEG, pupil response, heart and respiration rates, and salivary cortisol) and subjective stress ratings were collected. After collecting pre-stress neurophysiological activity in the first session, all participants underwent a cold pressor test (CPT) as an acute stressor. The second session was conducted immediately after the stress, and the third session was obtained to overlap with the expected peak of cortisol elevation. The fourth and fifth sessions were conducted 60 and 80 minutes, respectively, after the stress exposure.

The neurophysiological aspects of resilience have been studied using animal models, where individual animals are categorised into susceptible and resilient groups based on their display of depressive behaviour in response to stress^31-34^. In humans, the assessment of resilience is multifaceted and can include not only the observation of depressive behaviours following stress exposure but also mental activities that require a high degree of cognitive function, such as self-confidence and positive acceptance of the challenges faced^3,4^. Thus, although some aspects of resilience may be translated from animal models, exploring the neurophysiological responses underlying psychological resilience in humans can provide a more complete understanding of the mechanisms thereof.

Traditional studies have identified resilience signatures as individual differences in the intensity of well-known stress-related neurophysiological responses (stress-dependent model; Fig. 1c)^1,35^. However, spatially and temporally distinct resilience mechanisms may exist that cannot be observed using the conventional approach, which focuses only on known stress responses (stress-independent model; Fig. 1c). Personality questionnaires, which are not available for non-human animal studies, are suitable to directly explore the unknown neurophysiological dynamics involved in resilience. Given the relationship between neurophysiological responses and psychological questionnaires in the stress-dependent model, the strength of known neurophysiological stress responses could also explain individual differences in resilience, as assessed by psychological questionnaires. In contrast, in the stress-independent model, the neurophysiological responses to resilience explained by the questionnaire are independent of any known stress-related responses. To examine these two model, we assesseds acute stress-induced neurophysiological responses using a combination of peripheral and neuroimaging measurements, and examine the relationship between these responses and subjective assessments of psychological resilience.

## Results

We collected multiple non-invasive brain and peripheral responses before and after acute stress exposure (Fig. 1d) from 102 participants (77 males and 25 females). Individual resilience levels were measured using the Connor-Davidson Resilience Scale (CD-RISC) ^4^ (Supplementary Fig. 1). All participants underwent five brain imaging sessions and eight inter-scan-interval (ISI) sessions. At each imaging session, resting-state brain activity was recorded using functional magnetic resonance imaging (fMRI) and electroencephalography (EEG) to assess spatially- and temporally-resolved neural responses, as well as pupil size (Pup), electrocardiogram (ECG), and respiration (Resp). Each ISI included saliva samples for cortisol density (Cort) and subjective stress ratings (Subj). In addition to Pup, ECG, and Resp, a modified version of the CPT was administered as an acute stressor after the first brain imaging (baseline) session. All imaging data were evaluated as the change from the pre-stress baseline and labelled S2 to S5.

### Data do not support stress-dependent model

Our experiment successfully induced acute stress-related peripheral responses in the Pup, ECG, Cort, and Subj (Supplementary Fig. 2). Stress-related neural activity was also increased in resting-state fMRI (stress-driven functional connectivity [SFC] and Stress-related amygdala-insula connection [ScAI]) and resting-state EEG (Supplementary Fig. 3a-d, 4a-c; see Supplementary Discussion 1 for more details). To validate the stress-dependent resilience model (Fig. 1c), correlations between the CD-RISC scores and stress-related neurophysiological changes from baseline were verified. We found no statistically significant correlations in any peripheral responses through the experiments (Supplementary Fig. 2: *ρ* = -0.219∼0.186, *P_FDR_* = 0.320∼0.999) and no correlations in brain responses (Supplementary Fig. 3e,f, 4d,e: *r* = -0.143∼0.165, *P_FDR_* = 0.121∼0.958). These results indicated that our data did not support the stress-dependent model in resilience.

### Spatial features support the independent model

We explored novel whole-brain activities associated with individual differences in resilience. First, we evaluated the functional connectivity (FC) network changes that correlated with individual CD-RISC scores in post-stress four sessions (S2–S5). The number of resilience-related functional connections (RFC) was increased from S2 to S4 and then decreased at S5 (Fig. 2a,b). In S4, the two networks were significantly correlated with the CD-RISC scores (Fig. 2c,d; cluster-level *P_FDR_* < 0.05), both of which were negatively and positively correlated with the resilience score (RFCn and RFCp, respectively). The RFCn includes the bilateral insula, dorsal anterior cingulate (dACC), right orbitofrontal cortex (OFC), and right middle frontal gyrus (MFG), which overlap with the cortical salience network (SaN)^36^. RFCp included the right inferior frontal gyrus (IFG), precuneus (Prec), and posterior cingulate cortex (PCC), which overlapped with the cortical default mode network (DMN) (see Supplementary Fig. 5 for details). Importantly, the strength of correlations both in RFCn and RFCp gradually increased from S2 to S4 and attenuated in S5 (Fig. 2e): the coefficient in S4 was significantly higher than that in S2 (*T*_(87)_ = -2.787, *P_FDR_* = 0.020) and S5 (*T*_(87)_ = 3.04, *P*_FDR_ = 0.019) in RFCp^37^. Similar tendencies were also observed in RFCn, but did not reach statistical significance (S2 and S4: *T*_(87)_ = 2.095, *P_FDR_* = 0.10; S4 and S5: *T*_(87)_ = -2.486, *P*_FDR_ = 0.089) (see Supplementary Fig. 6 for another resilience-related functional connections, identified in S2 [RFCs2]).

**Fig. 2:**
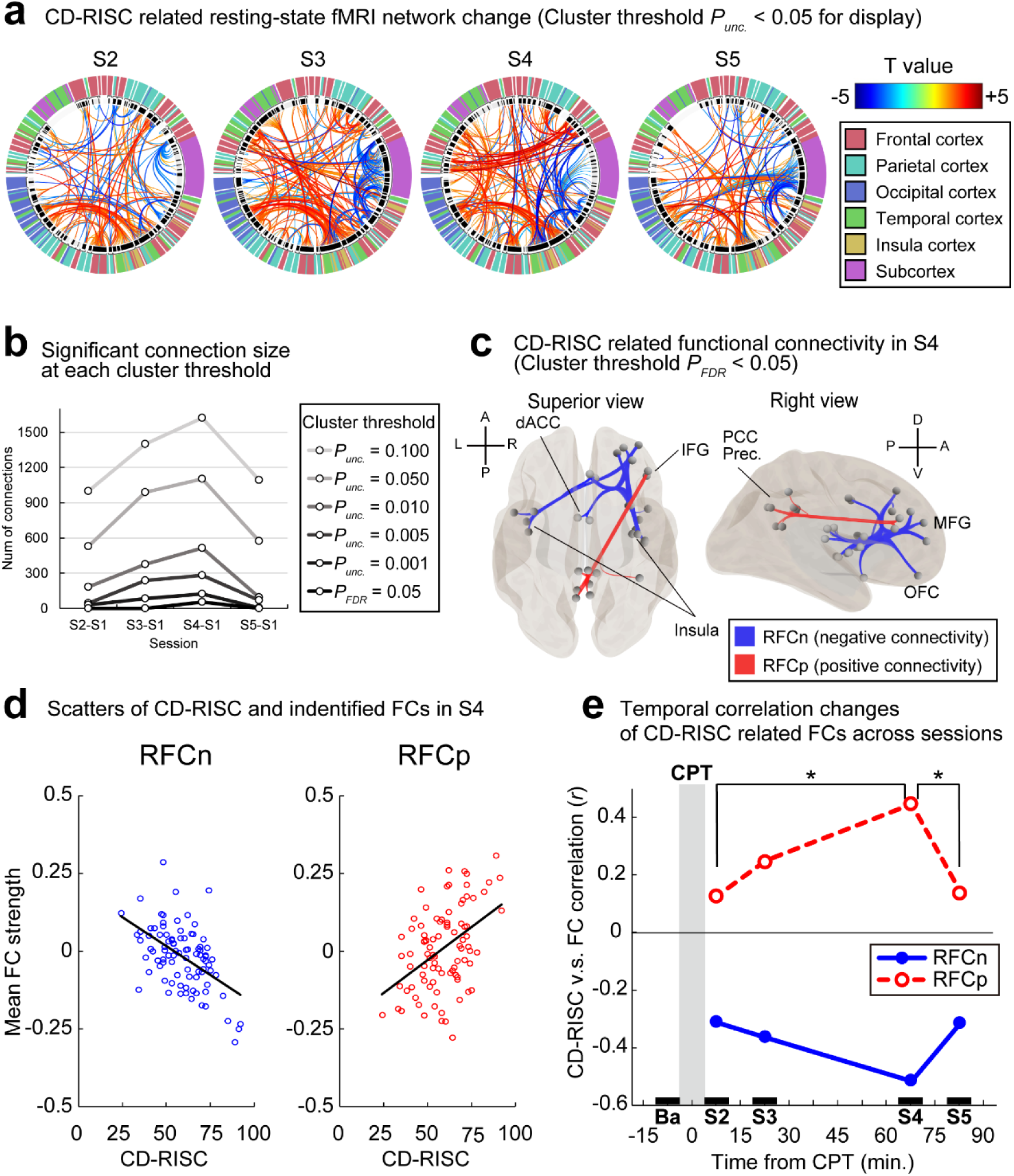
Correlated fMRI functional connectivity with resilience peaked 60 minutes after the acute stress experience. Change in the number of significant connections as a function of time. (**a**) The connectome ring depicts the number of CD-RISC-related functional connectivity (FC) from S2 to S5. The statistical threshold for the ring was set at *P* < 0.05, uncorrected, for display purposes. The number of FCs increased as a function of time, peaking at S4 (approximately 60–70 min after acute stress) and then decreasing at S5 (approximately 75–85 min after acute stress). (**b**) The number of significant FCs was greatest in S4, regardless of the statistical thresholds (both uncorrected and corrected). (**c**) Two significant FCs were identified at S4 with a cluster-level threshold *P_FDR_* of < 0.05. FC negatively correlated with CD-RISC scores (RFCn; shown as blue lines) in the bilateral insula, dorsal anterior cingulate (dACC), right orbitofrontal cortex (OFC), and right middle frontal gyrus (MFG). Another FC positively correlated with CD-RISC scores (RFCp; shown in red lines), including the right inferior frontal gyrus (IFG), precuneus (Prec.), and posterior cingulate cortex (PCC). The specific ROI names and locations in the brain are shown in Supplementary Fig. 5. (**d**) Scatter plots of individual CD-RISC scores and FC strengths in RFCn (*r* = -0.517, *P_FDR_* = 1.84E-7) and RFCp (*r* = 0.447, *P_FDR_* = 1.01E-5) at S4. (**e)** Temporal dynamics of the correlation between CD-RISC and FCs in RFCn and RFCp. The correlation coefficient (*r*) of RFCp was significantly higher in S4 than in S2 and S5. \**P_FDR_* < 0.05, \*\**P_FDR_* < 0.01, corrected.

Previous non-human animal studies have primarily highlighted the contribution of subcortical regions^38^. To evaluate the contribution of the subcortex to psychological resilience, we calculated the spontaneous neural activity using the fractional amplitude of low-frequency fluctuations (fALFF). Based on a review of prior animal resilience studies^38^, seven subcortical regions were selected as the regions of interest (ROIs) (Supplementary Fig. 7). In S4, fALFF magnitudes in the bilateral posterior hippocampus (HIPp) positively correlated with individual CD-RISC scores (Supplementary Fig. 7a,b: *r*_(90)_ = 0.296, *P_FDR_* = 0.033). In the other six ROIs, this analysis failed to detect significant correlations with CD-RISC (Supplementary Fig. 7c).

### Temporal features support the independent model

In animal models, individual differences in resilience are represented by differences in brain rhythms, measured using local field potentials (LFPs)^34,39^. Similar differences are probably expressed in the human brain as well. We, therefore, evaluated the spontaneous human brain rhythms using EEG (Supplementary Fig. 8). In S4, we identified two frequency components that peaked at 26.5 Hz (E26.5 in beta 2 band) and 43.0 Hz (E43.0 in gamma band), which were negatively correlated with individual CD-RISC scores (Fig. 3b,d, *P_FDR_* < 0.05). Both components were spatially distributed from the right frontal region to the bilateral parieto-occipital region in S4. This negative correlation partially remained during S5, but only in the parieto-occipital regions at E26.5 (Fig. 3b, in S5, *P_FDR_* < 0.05). The temporal correlation changes highlighted that the correlation was not observed during S2 (E26.5: *r*_(88)_= -0.091, *P_FDR_* = 0.535, E43.0: *r*_(88)_= -0.116, *P_FDR_* = 0.374) or S3 (E26.5: *r*_(88)_= 0.007, *P_FDR_* = 0.951, E43.0: *r*_(88)_= -0.060, *P_FDR_ =* 0.578), instead appearing >60 minutes later in S4 (E26.5: *r*_(88)_= -0.395, *P_FDR_* = 0.001, E43.0: *r*_(88)_= -0.369, *P_FDR_* = 0.002) and were sustained till S5 (E26.5: *r*_(88)_= - 0.304, *P_FDR_* = 0.008, E43.0: *r*_(88)_= -0.255, *P_FDR_* = 0.033). The correlation coefficient was stronger in S4 than in S2 (E26.5: *T*_(87)_ = 2.942, *P_FDR_* = 0.013; E43.0: *T*_(87)_ = 2.745, *P_FDR_* = 0.022) and S3 (E26.5: *T*_(87)_ = 3.698, *P_FDR_* = 0.002; E43.0: *T*_(87)_ = 3.446, *P_FDR_* = 0.005), while the coefficient in S5 was stronger than in S2 (*T*_(88)_ = 2.760, *P_FDR_* = 0.014) only in E26.5 (Fig. 3d). We confirmed that the identified activations were not correlated with other stress-related measures (Supplementary Fig. 9), but were only correlated with the psychological resilience score, CD-RISC.

**Fig. 3:**
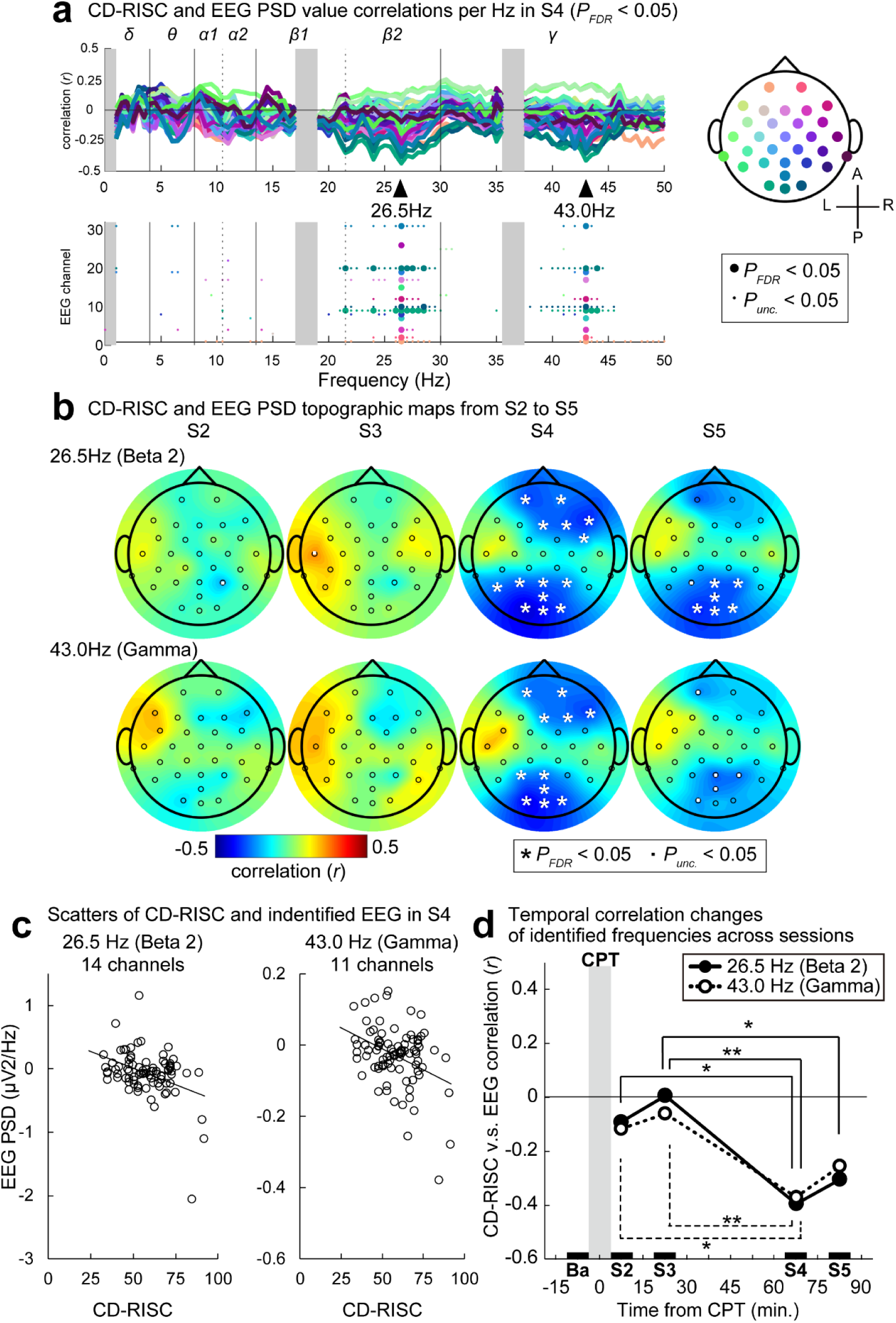
Spectral EEG features correlated with resilience 60 min after the acute stress experience. (**a**) Top: The correlation coefficients between the individual CD-RISC scores and the power spectral density (PSD) for each frequency band at S4. The colour of each line corresponds to the 32 channels located on the scalp. Correlation peaks were identified at 26.5 and 43.0 Hz. Bottom: The statistically significant frequencies in each channel. Large and small dots indicate a significant channel and frequency at the threshold of *P_FDR_* < 0.05 corrected and *P_unc._* < 0.05, respectively. Frequency bands shown in grey masks are not analysed due to noise from the fMRI scanner. See Supplementary Fig. 8 for correlations between CD-RISC and EEG in other sessions. (**b**) Temporal changes of topographic maps from S2 to S5 at 26.5 (Beta 2) and 43.0 (Gamma) Hz. White asterisks indicate statistically significant EEG channels with *P_FDR_* < 0.05, corrected. (**c**) Scatter plots of individual CD-RISC scores and means of PSDs taken from statistically significant channels at 26.5 (*r* = -0.395, *p* = 1.55E-4) and 43.0 Hz (*r* = -0.304, *p* = 0.004) in S4. (**d**) Temporal dynamics of the correlation between CD-RISC and PSD at 26.5 and 43.0 Hz. The correlation coefficient (*r*) of 26.5 Hz was significantly lower in S4 than in S2 and S3, and lower in S5 than in S3. The coefficient of 43.0 Hz was significantly lower in S4 than in S2 and S3. \**P_FDR_* < 0.05, \*\**P_FDR_* < 0.01, corrected.

### Impact level of identified factors

We tested 13 neural and peripheral factors at four time points (13 × 4 = 52) that were potentially related to individual differences in resilience. However, the critical question remains as to whether individual resilience can be predicted. Thus, the elastic net was used to calculate the best combination of factors and their impact on psychological resilience. The elastic net optimisation process from the 52 factors revealed a model that included six factors (Fig. 4a, Model 1: adjusted *R^2^* = 0.322, *P* = 1.07E-05). Based on this optimised model, we found that the most contributed factor was RFCn (standardised partial regression coefficient β = -0.254, contribution level [CL] = 40.60%); the second factor was RFCp in S4 (*β* = 0.204, CL = 32.57%); factors 3–5 were E43.0 (*β* = -0.056, CL = 8.89%), HIPp (*β* = 0.052, CL = 8.31%), and E26.5 (*β* = -0.047, CL = 7.48%) in S4, respectively (Fig. 4b). This analysis showed that the stress-driven brain or peripheral responses reported in previous studies showed negligible contribution to predict individual psychological resilience in humans (see Supplementary Fig. 10 and 11 for the robustness of our results).

**Fig. 4:**
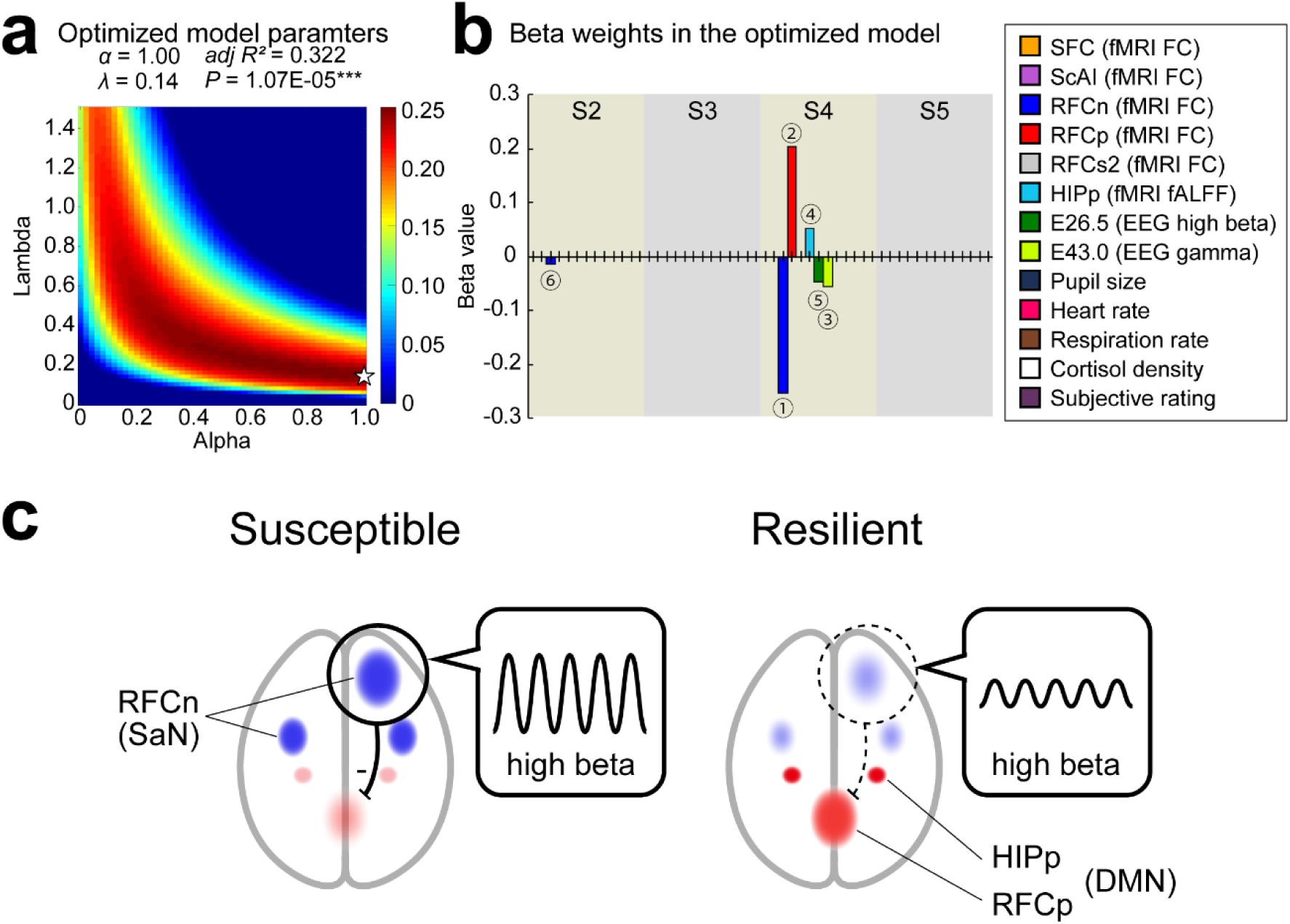
Important neural factors for psychological resilience and a proposed neural model of human psychological resilience. (**a**) The heat map shows the R^2^ values during the process of optimising the alpha and lambda parameters for reguralisation term using the elastic net. The star in the map indicates the peak value. This optimisation yielded the best six factors to predict individual resilience (Model 1: *α* = 1.00, *λ* = 0.14, adjusted *R^2^* = 0.322, *P* = 1.07E-05) out of 52 (13 measurements × 4 sessions) factors. (**b**) The weights of the six selected factors indicate that RFCn, which overlaps with SaN, and RFCp, which overlaps with DMN at S4, contribute dominantly to the prediction of individual resilience levels. The circled numbers indicate the ascending order of the absolute value of the optimised weights. (**c**) Model of the neural mechanism of human resilience based on the current findings. In this model, sustained activation of the middle frontal gyrus, part of the SaN, suppresses activation of the medial parietal lobe, part of the DMN, through high-beta oscillations. This over-suppression occurs approximately 60 minutes after stress forms susceptible individuals, and the disinhibition of the DMN contributes to forming resilient individuals.

## Discussion

We applied a multimodal approach to elucidate the neurophysiological characteristics of human resilience and observed that their individual differences peaked 60 min after stress exposure. The identified neural characteristics are independent of neurophysiological responses focused on stress-dependent activities that end within 60 min, and are potential therapeutic targets that effectively leverage individual resilience. Although previous human studies have examined the correlation between individual resilience scores and resting-state brain activity^38,40^, we provide several advancements beyond the knowledge of resilience from prior animal models and existing imaging studies. First, most animal studies have focused on individual differences in the responses to stress exposure, whereas most human studies have not experimentally introduced stress exposure to participants. In contrast, we directly observed the neurophysiological responses to actual stress. Second, previous studies with acute stressors have not focused on responses longer than 1 hour because autonomic and hormonal responses apparently subside within 1 hour. However, physiological responses persist even after 1 hour of stress exposure^25-27^, and the neural states more than 1 hour after stress are significantly different from the pre-stressed state^28-30^. We, therefore, examined neurophysiological dynamics for more than 60 min, which has been overlooked in conventional resilience studies.

Accordingly, we identified six factors correlated with individual resilience. Furthermore, the machine learning approach revealed that the most influential factors were RFCn and RFCp, both from the FC in S4. This result indicates that human resilience is supported by the SaN and DMN in the neocortex, rather than the subcortex system, which has been the focus for many years. In particular, the insular cortex is potentially a core region in determining individual resilience and susceptibility after acute stress because the insula is thought to be the hub of the SaN and suppresses the posterior part of the DMN^41^. A further discussion of each identified factor is provided in Supplementary Discussion 2.

The combination of fMRI and EEG revealed that resilience-related EEG changes in the prefrontal area were not only observed simultaneously, but also had a spatially similar distribution to the FC increases observed in fMRI. Particularly, increases in RFCn (SaN) and E26.5 (high-beta) in people with lower resilience may capture the same underlying neural machinery. Indeed, a study investigating the resting-state natural frequency reported that high-beta activity (23.3∼27.1 Hz) is dominant in the middle frontal gyrus (MFG)^42^. In conjunction with this report, in individuals with lower resilience, the dominance of high-beta power in the MFG may be enhanced by activation of the SaN (Fig. 4c). In contrast, an increase in the DMN in the medial parietal lobe and a decrease in high-beta power in the parietal channels was observed in resilient individuals. A similar phenomenon of reduced high-beta power and increased DMN activity has been reported^43^, and it is possible that high-beta activity suppresses DMN activity in the parietal and occipital regions, especially in lower resilient individuals.

Our results also showed that the increase in high-beta power was widely distributed from the prefrontal to the parieto-occipital regions, and that high-beta activity potentially played an important role in the suppression of the DMN by the SaN (Fig. 4C). To test this hypothesis, it is necessary to artificially suppress high-beta activity after stress. The next research step would be to test the neuromodulation effects, whether the suppression of high-beta activity (1) reduces the activation of the SaN in the prefrontal cortex while enhancing that of the DMN in the parietal cortex, and (2) contributes to individual resilience levels in real life.

Our study highlights the specific time window that drives neurophysiological resilience. Considering that the physical stress used in the present study is only one aspect of the wide range of stressors that we encounter, we need to explore such a time window for other stressors. Targeting interventions at such specific time windows could maximise the effectiveness of protection and treatment of acute stress-related disorders such as post-traumatic stress disorder (PTSD).

## Methods

### Participants

All experiments were conducted in accordance with the principles of the Declaration of Helsinki and were approved by the ethical review board (approval number 177) of the Kochi University of Technology. In total, 102 participants (77 males; mean age 20.11 [range 18–25] years) with normal or corrected-to-normal vision participated in this study. Eight participants were left-handed.

The sample size of the participants was predetermined based on the size of recent psychological resilience-related studies using resting-state fMRI data^44-46^. All participants provided informed consent prior to the experiments and completed a resilience-related personality trait test twice on different days. Four individuals with unstable scores (change >20 points) on the resilience questionnaire were excluded from the analysis. Two participants dropped out of the study.

Data from the remaining 96 participants were used to analyse the physiological data (72 males; mean age, 20.13 years). For the brain imaging analyses, we excluded three individuals with brain morphological abnormalities and two individuals who were prescribed psychiatric medications. During the experiment, we failed to collect one fMRI and two EEG data sets because of machine problems. Therefore, we analysed data from the remaining 90 participants for fMRI (68 males; mean age, 20.13 years) and 88 participants for EEG (67 males; mean age, 20.17 years).

### Experimental procedures

#### Participant experiences

All neurophysiological experiments were conducted between 12:00 and 20:00 to minimise the diurnal effects of cortisol levels. After completing the four questionnaires, an electrocardiogram (ECG), a respiratory belt transducer (Resp), and an electroencephalogram (EEG) were attached to all participants. The experiment consisted of five brain imaging sessions with eight physiological measurements between sessions (Fig. 1d). During each brain imaging session, resting-state fMRI and EEG data were collected concurrently for 10 min. During the session, the participants were required to open their eyes and fixate on a cross displayed in the centre of the screen through a mirror in the scanner. The pupil size (Pup), ECG, and Resp data were collected during each session. At the intersession interval (ISI), saliva samples were collected, and participants reported their current feelings by moving a slider displayed on the screen. Pup, ECG, and Resp information were collected during this interval. Each interval lasted for 2 minutes.

Between brain imaging sessions 1 and 2, all participants underwent a 2-minute CPT as an acute stressor in the MRI scanner. Between sessions 3 and 4, the participants took breaks outside the scanner, during which they were informed that no further stress would be applied during the experiment.

#### Questionnaires

All participants completed four questionnaires twice: a couple of days before and on the day of the experiment, approximately 40 min before entering the MRI scanner. The main questionnaire used in this study was the Connor-Davidson Resilience Scale (CD-RISC), which assesses individual levels of resilience to stress^4^. This scale is considered the most dominant to assess resilience, as it has the best psychometric properties^47,48^. It comprises 25 questions scored from 0 to 100. Individuals with a higher score are more resilient. In addition, we used the Stress Mindset Measure (SMM)^49^ to assess individual ‘stress-is-enhancing’ or ‘stress-is-debilitating’ mindsets, the Perceived Stress Scale (PSS)^50^ to assess chronic daily stress levels, and the Beck Depression Inventory-II (BDI)^51^ to assess individual depression levels. All questionnaires were translated into Japanese and the participants completed the forms using their smartphones or personal computers.

#### Acute stress manipulation

Participants underwent a modified version of the CPT, essentially mimicking previous studies^18,19^, but used an ice glove instead of ice water as the stressor to avoid the risk of spilling water in the scanner. The gloves were chilled to approximately -20°C to match the subjective stress level of placing the hands in ice water. After instructions regarding the stress procedure, the experimenter placed an ice glove on the participant’s left hand for 2 min to induce a physiological stress response. Throughout the procedure, participants maintained a supine position in the scanner and fixated on a central cross displayed on the screen. Participants were able to stop the stress procedure in the middle of the test by pressing an emergency bulb in their right hand. The participants were informed that their faces would be recorded using a video camera. The 2-minute duration of our procedure differed from the 3-minute duration typically used for CPT^18,19^. The decision to use a shorter duration was based on previous studies showing that, in some experiments, more than 20% of participants are unable to keep their hands in cold water beyond the 2-minute mark^52-54^. Despite a shorter exposure duration, this procedure was effective (Supplementary Fig. 2d). EEG, pupil size, ECG, Resp, and saliva samples were collected during the CPT. Participants also reported their subjective feelings immediately after the test.

### Equipment for data acquisitions

#### Software

We used MATLAB R2019a with Psychtoolbox 3.0.12 for stimulus presentation and data analysis. In addition, SPSS ver. 23 (IBM Corp. Armonk, NY) and R-studio ver. 2022.07.2 were used for the statistical tests and machine learning approaches.

#### fMRI and EEG

A 3T MRI scanner (Siemens Prisma, Munchen Germany) with a 64-channel head coil was used. Functional images were acquired using an echo-planar imaging (EPI) sequence (repetition time [TR]: 2.0 s; echo time [TE]: 27 ms; flip angle [FA]: 90°; 36 slices; slice thickness: 3 mm; in-plane resolution: 2 × 2 mm; descending). We did not use multiband accelerated imaging because the guidelines for simultaneous fMRI-EEG recordings provided by Brain EEG Products (Germany) did not permit it. Each functional scan was comprised of 300 scans. High-resolution anatomical images were acquired using an MP-RAGE T1-weighted sequence (TR: 2,400 ms; TE = 2.32 ms; FA: 8°; 192 slices; slice 478 thickness: 0.9 mm; in-plane resolution: 0.93 × 0.93 mm). The anatomical image was acquired twice, before sessions 1 and 5, to account for changes in the head position after the break outside the scanner.

EEG data were acquired from MRI-compatible 32-sintered Ag/AgCl electrodes with extended scalp coverage, including 31 scalp recording electrodes and one electrode on the upper back for ECG (BrainCap-MR; Brain Products, Bayern Germany). Electrodes AFz and FCz served as ground and online references, respectively. All signals were recorded with a bandpass of 0.016–250 Hz and were digitised at 5,000 Hz (BrainAmp MR Plus, Brain Products). Electrode impedances were lowered to <10 kΩ prior to recording and monitored throughout the experiment.

### Peripheral physiological measurements

#### Pupil size

Eye movements and pupil size were recorded using EyeLink 1000 PLUS (SR Research Ltd., Ottawa Canada). The pupil diameter was tracked from the right eye at a sampling rate of 500 Hz, measured in centroid mode, to reduce noise in the pupil data. This system was also used to confirm whether the participants were awake.

#### Heart rate and respiration

An MRI-compatible three-lead ECG and contractible belt (BIOPAC Systems Inc. Goleta, CA) were attached to the chest and abdomen, respectively, to detect the timing of the R peaks and respiratory rhythm. The sampling rate was 10,000 Hz.

#### Saliva

To obtain cortisol data, saliva samples were collected using a SalivaBio Oral Swab (Salimetrics, LLC, Carlsbad, CA). Participants were required to put the swab under their tongue for 2 min. The collected swab was placed in a single centrifuge tube and frozen in cold storage at -50°C.

#### Subjective ratings

The participants rated their momentary stress levels eight times during the experiment, in the scanner (seven times) or outside the scanner (once during the break). For each rating, one question, “How stressed are you feeling right now?” was displayed on the screen and they answered it on a scale from -50 to 50 (by moving the slider from 0). In addition, immediately after the CPT, participants rated their subjective feeling of unpleasantness and pain levels following the stress rating.

### Statistical analyses

#### Resting-state fMRI

Preprocessing procedures and statistical analyses were performed using the CONN Functional Connectivity Toolbox (ver. 20.b: www.nitrc.org/projects/conn), and SPM12 (www.fil.ion.ucl.ac.uk/spm/). The default preprocessing pipeline for volume-based analysis provided by CONN was used. For each participant, the EPI images were temporally realigned across volumes and sessions, and each slice was time-shifted and resampled using interpolation to match the middle of each acquisition time (slice-timing correction). Artefact detection tool (ART)-based scrubbing was performed to detect outliers. The outlier detection threshold was set at the 95th percentile for the normative samples. Functional and anatomical data were normalised to the standard Montreal Neurological Institute space and segmented into grey matter, white matter, and cerebrospinal fluid (CSF) tissue classes using the SPM12 unified segmentation and normalisation procedure. Finally, the images were smoothed using a Gaussian kernel of 8 mm full width at half maximum. Before the first level analysis, denoising was performed. Linear trends, low-frequency drift, and high-frequency noise in the blood-oxygen-level dependent (BOLD) signal were removed using a temporal bandpass filter (0.008∼0.09 Hz) to focus on low-frequency fluctuations. Noise components from the white matter (five components), cerebrospinal areas (five components), estimated subject motion parameters (12 components), estimated outlier scans, and session effects were removed by regression analysis. Additionally, to denoise peripheral cardiac and respiration artefacts, the collected ECG and Resp data were modelled using RETROICOR with the PhysIO toolbox^55,56^ and were applied as noise regressors (20 components).

#### ROI-to-ROI analysis

We used an original set of ROIs, with 360 regions in the cerebral cortex defined based on Glasser’s template^57^, 54 ROIs in the subcortex defined based on Tian’s template^58^, and 12 ROIs in the brainstem defined based on the AAL3^59^. In total, 426 ROIs were used for the FC analysis. The cerebellum was excluded from ROI-based analysis. Individual correlation maps were generated by extracting the mean BOLD time course from each ROI and calculating the correlation coefficients between two ROIs. We compared the correlation maps across sessions; therefore, the ROI order was optimised using baseline session one. This order was defined using a data-driven hierarchical clustering procedure (complete-linkage clustering^60^) implemented in CONN. Group-level contrasts were identified as changes from the baseline session and reported after subtracting the effects of sex and age regressors. Parametric statistics based on functional network connectivity were used to report the results^61^. A cluster-level FDR-corrected p-value < 0.05 was applied to report the significant FC networks^62^.

#### Seed-to-voxel analysis

To confirm consistency with previous stress-induced FC changes reported by van Marle et al.^20^, we applied seed-based correlation analysis. The bilateral amygdala was used as the seed region. In accordance with the previous report, the statistical threshold was set at *P* < 0.001, uncorrected.

#### fALFF

fALFF maps represent a relative measure of BOLD signal power within the frequency band of interest (e.g., 0.01–0.10 Hz) compared to that over the entire frequency spectrum. We used these to assess the BOLD changes in specific subcortical regions reported as resilience-related systems^38^. Correlations between individual CD-RISC scores and the bilateral anterior hippocampus, posterior hippocampus (HIPp), amygdala, nucleus accumbens, ventral tegmental area, locus coeruleus and dorsal raphe nucleus were evaluated. The amplitudes of the left and right ROI were averaged. The significance of the correlation was corrected using an FDR *P* < 0.05.

#### Data collection from Neurosynth datasets

Neurosynth is a meta-analysis tool that displays brain activity maps corresponding to cognitive functions by analysing large amounts of previously accumulated fMRI data^63^. This tool is useful to avoid inappropriate inferences regarding the relationship between brain activity and cognitive function. As shown in Supplementary Fig. 3C, this tool was used to examine whether the seeds of the stress-related functional networks identified in our analysis overlapped with previously reported stress-related brain activity. The word ‘stress’ was used to search for brain activity, and a brain activity map (uniformity test *P* < 0.01, FDR-corrected) based on 321 studies is shown.

### EEG

#### Resting state

Preprocessing procedures and statistical analyses were performed using a Brain Vision Analyser 2 (BVA2; Brain Products) and EEGLAB (ver. 2021.1, https://sccn.ucsd.edu/). First, in BVA2, the MR gradient artefact was removed using an average template subtraction method^64^, and the resulting EEG data were down-sampled to 500 Hz.

Second, the EEG data were bandpass-filtered between 1∼200 Hz. Third, cardiovascular artefacts were removed using data from one ECG channel. Denoising was performed using a semiautomatic template-matching procedure. Fourth, eyeblink correction was performed using an independent component analysis (ICA). Because specific sensors for vertical/horizontal electrooculography (EOG) were not attached to the scanner for safety reasons, we used the EEG data obtained from electrode FP1 as a proxy for the vertical activity of the EOG channel.

Further denoising and analysis were performed using custom scripts and EEGLAB, as follows. The 10 min continuous EEG data were segmented into 300 epochs with 2.0 sec duration, time-rocked to the TR of the fMRI. To detect abnormally noisy channels due to insufficient adhesion between the scalp and electrode, the power spectrum density (PSD) distribution of the channel with the highest signal intensity was compared with the average PSD distribution of the other top ten PSD intensity channels. If PSD distribution in the given channels was significantly different from the average of the other top 10 channels (Wilcoxon rank-sum test), the channel was excluded, and the data were interpolated by the corresponding channel by replacing it with the averaged EEG data from the neighbouring channels. Thereafter, the noise in each epoch was inspected and rejected with the following parameters: abnormal values, over ±120 μV; abnormal trends, 50 μV/epoch; R^2^ limit, 0.3; improbable data, 5 std.; abnormal distribution, 5 std.; abnormal spectra (slow), over -60∼40 dB, 0∼40 Hz. On average, 12.10 ± 9.65% of the epochs were excluded with these criteria, Finally, we applied ICA-based denoising techniques implemented in EEGLAB (‘runica’ function). Each independent component was labelled using the ICLabel plugin (https://github.com/sccn/ICLabel). The independent components with < 95% probability of the brain signal were flagged as IC artefacts and rejected. For EEG frequency analysis, each epoch data was Fourier transformed to PSD with a frequency resolution of 0.5 Hz, and deleted epochs were filled by the spline interpolation method. We excluded 0–0.5, 17.5–18.5, and 36.0–37.0 Hz from the PSD data because TR-driven fMRI noise components remained dominant even after several noise reduction procedures. The frequency bands were divided into delta (0.5∼3.5 Hz), theta (4.0∼7.5 Hz), alpha1 (8.0∼10.0 Hz), alpha2 (10.5∼12.5 Hz), beta1 (13.0∼21.5 Hz), beta2 (21.5∼29.5 Hz), and gamma (30.0∼49.5 Hz).

### Peripheral physiological data

#### Pupil size

The mean diameter (µm) during each brain imaging session (10 min), ISI (2 min), and CPT (2 min) was used as a representative value for each participant. Some data were lost because of partially open eyes or eyes covered with eyelashes. Data with <70% acquisition per session were excluded. The probability of the remaining valid data was 89.0% for the total sample. Spearman correlation coefficients were calculated between the pupil diameter and individual CD-RISC scores at each time point.

#### ECG analysis

Beats per minute (BPM) were calculated from R-peak information. The data were down-sampled to 1,000 Hz for the analysis. The ECG data contained noise owing to electromagnetic induction derived from rapid magnetic field changes during the MRI scan. A customised Pan–Tompkins ECG QRS detector^65,66^ was used to detect the R–R intervals from the data obtained during scanning. Almost all R peaks were detected automatically by this algorithm, but some data were manually detected by the experimenter. The probability of the remaining valid data was 98.3% of the total sample. Spearman’s correlation coefficients between the BPM and individual CD-RISC scores were calculated at each time point. In addition, although we attempted to calculate heart rate variability from the R peaks, this was abandoned because it is often difficult to detect accurate R-peak positions owing to MRI-derived noise.

#### Respiratory rate

In contrast to the ECG data, the respiratory data were free of MRI-derived noise because they measured changes in air pressure in a belt worn over the abdomen. Data were down-sampled to 1,000 Hz for analysis. Exhalation and inhalation peaks were detected, and respirations per minute (RPM) were calculated and averaged for each session. The probability of valid data was 99.6% for all the samples. Spearman’s correlation coefficients were calculated between the RPM and individual CD-RISC scores at each time point.

#### Cortisol concentration

Saliva samples were analysed in the laboratory using a salivary cortisol enzyme-linked immunosorbent assay (ELISA) kit (Salimetrics). The cortisol concentrations (µg/dL) during each ISI (2 min), CPT (2 min), and break outside the scanner (2 min for sampling) were evaluated. Although almost all saliva samples were analysed in duplicate and the average optical density was calculated to reduce measurement error, some collected samples were tested once, and the other samples were not tested owing to insufficient sample quantity. The probability of the remaining data being valid was 83.9% for the total sample. Spearman’s correlation coefficients were calculated for cortisol density and individual CD-RISC scores at each time point.

### Machine learning

We used linear regression models with a machine learning approach to evaluate the best combinations to predict individual resilience levels from the observed neurophysiological data. The best parameters were selected by the elastic net, which regularised the L1 and L2 norms. We validated and reported eight neural factors (SFC, ScAI, RFCn, RFCp, RFCs2, HIPp, E23.5, and E43.0), four peripheral physiological factors (Pup, BPM, Resp, and Cort), and one subjective rating (Subj) potentially related to individual stress or resilience responses. Thirteen modalities at four time points (52 factors) were selected as explanatory factors. All values were subtracted from the value of the baseline session one and z-normalised before estimation.

Missing data for explanatory variables were interpolated using the mean values of the remaining participants. In the main model, the individual CD-RISC scores (n = 90) were calculated using 52 parameters. In the second model, 13 modalities were used, but were optimised per session (S2–S5). In the third model, only six neural modalities (RFCn, RFCp, RFCs2, HIPp, E23.5, and E43.0) showing significant correlations were used as explanatory factors. Alpha and Lambda hyperparameters were optimised by nine-fold cross-validation. The search range of alpha was 0–1.0 per 0.02, and that of lambda was 0–1.5 per 0.02.

### Data and code availability

The data that support the findings of this study are available on OSF: https://osf.io/kvnh9/

## Acknowledgements

We would like to thank M. R. Delgado at Rutgers University for comments on the manuscripts, K. Nakahara, K. Jimura, H. Kadota, M. Iwata in Kochi University of Technology for technical support and useful discussion, and T. Furuyashiki at Kobe University and K. Nakamura at Kansai Medical University for useful comments and discussions. This work was financed by KAKENHI from Japan Society for the Promotion of Science (JP21K07262, JP21H00211, and JP22H05219 to N.W., JP20H00521 and JP21K18267 to M.T., JP22K12786 to S.Y.), a grant from Public Health Research Foundation, a grant from The Naito Science & Engineering Foundation to N.W., and a grant from Takeda Science Foundation to M.T.

## Author contributions

N.W. and M.T. conceived the project. N.W. and R.K. collected data. N.W. and S.Y. analysed data. M.T. directed and advised on the data analysis. N.W. and M.T. wrote the manuscript.

## Corresponding author

Correspondence to N.W. or M.T.

## Additional information

None

## References

1 Feder, A., Nestler, E. J. & Charney, D. S. Psychobiology and molecular genetics of resilience. Nature Reviews Neuroscience 10, 446–457 (2009).

2 Fletcher, D. & Sarkar, M. Psychological resilience. European psychologist (2013).

3 Southwick, S. M., Vythilingam, M. & Charney, D. S. The psychobiology of depression and resilience to stress: implications for prevention and treatment. Annu. Rev. Clin. Psychol. 1, 255–291 (2005).

4 Connor, K. M. & Davidson, J. R. Development of a new resilience scale: The Connor-Davidson resilience scale (CD-RISC). Depression and anxiety 18, 76–82 (2003).

5 Bonanno, G. A. Loss, trauma, and human resilience: Have we underestimated the human capacity to thrive after extremely aversive events? American psychologist 59, 20 (2004).

6 Bradley, M. M., Miccoli, L., Escrig, M. A. & Lang, P. J. The pupil as a measure of emotional arousal and autonomic activation. Psychophysiology 45, 602–607 (2008).

7 De Berker, A. O. et al. Computations of uncertainty mediate acute stress responses in humans. Nature communications 7, 10996 (2016).

8 Kudielka, B. M., Buske-Kirschbaum, A., Hellhammer, D. H. & Kirschbaum, C. Differential heart rate reactivity and recovery after psychosocial stress (TSST) in healthy children, younger adults, and elderly adults: the impact of age and gender. International journal of behavioral medicine 11, 116–121 (2004).

9 Schiweck, C., Piette, D., Berckmans, D., Claes, S. & Vrieze, E. Heart rate and high frequency heart rate variability during stress as biomarker for clinical depression. A systematic review. Psychological medicine 49, 200–211 (2019).

10 Yackle, K. et al. Breathing control center neurons that promote arousal in mice. Science 355, 1411–1415 (2017).

11 Boiten, F. A., Frijda, N. H. & Wientjes, C. J. Emotions and respiratory patterns: review and critical analysis. International Journal of Psychophysiology 17, 103–128 (1994).

12 Hermans, E. J. et al. Stress-related noradrenergic activity prompts large-scale neural network reconfiguration. Science 334, 1151–1153 (2011).

13 Chang, P. F., Arendt-Nielsen, L. & Chen, A. C. Dynamic changes and spatial correlation of EEG activities during cold pressor test in man. Brain research bulletin 57, 667–675 (2002).

14 Shao, S., Shen, K., Yu, K., Wilder-Smith, E. P. & Li, X. Frequency-domain EEG source analysis for acute tonic cold pain perception. Clinical Neurophysiology 123, 2042–2049 (2012).

15 Stevens, A., Batra, A., Kötter, I., Bartels, M. & Schwarz, J. Both pain and EEG response to cold pressor stimulation occurs faster in fibromyalgia patients than in control subjects. Psychiatry research 97, 237–247 (2000).

16 Kirschbaum, C., Pirke, K.-M. & Hellhammer, D. H. The ‘Trier Social Stress Test’– a tool for investigating psychobiological stress responses in a laboratory setting. Neuropsychobiology 28, 76–81 (1993).

17 Goodman, W. K., Janson, J. & Wolf, J. M. Meta-analytical assessment of the effects of protocol variations on cortisol responses to the Trier Social Stress Test. Psychoneuroendocrinology 80, 26–35 (2017).

18 Schwabe, L., Haddad, L. & Schachinger, H. HPA axis activation by a socially evaluated cold-pressor test. Psychoneuroendocrinology 33, 890–895 (2008).

19 Schwabe, L. & Schächinger, H. Ten years of research with the Socially Evaluated Cold Pressor Test: Data from the past and guidelines for the future. Psychoneuroendocrinology 92, 155–161 (2018).

20 Van Marle, H. J., Hermans, E. J., Qin, S. & Fernández, G. Enhanced resting-state connectivity of amygdala in the immediate aftermath of acute psychological stress. Neuroimage 53, 348–354 (2010).

21 Zhang, W. et al. Acute stress alters the ‘default’ brain processing. Neuroimage 189, 870–877 (2019).

22 Tops, M., van Peer, J. M., Wester, A. E., Wijers, A. A. & Korf, J. State-dependent regulation of cortical activity by cortisol: an EEG study. Neuroscience letters 404, 39–43 (2006).

23 van Peer, J. M., Roelofs, K. & Spinhoven, P. Cortisol administration enhances the coupling of midfrontal delta and beta oscillations. International Journal of Psychophysiology 67, 144–150 (2008).

24 Quaedflieg, C., Meyer, T., Smulders, F. & Smeets, T. The functional role of individual-alpha based frontal asymmetry in stress responding. Biological psychology 104, 75–81 (2015).

25 Steptoe, A., Hamer, M. & Chida, Y. The effects of acute psychological stress on circulating inflammatory factors in humans: a review and meta-analysis. Brain, behavior, and immunity 21, 901–912 (2007).

26 Slavish, D. C., Graham-Engeland, J. E., Smyth, J. M. & Engeland, C. G. Salivary markers of inflammation in response to acute stress. Brain, behavior, and immunity 44, 253–269 (2015).

27 Joëls, M. Corticosteroids and the brain. Journal of Endocrinology 238, R121–R130 (2018).

28 Hermans, E. J., Henckens, M. J., Joëls, M. & Fernández, G. Dynamic adaptation of large-scale brain networks in response to acute stressors. Trends in neurosciences 37, 304–314 (2014).

29 Quaedflieg, C. et al. Temporal dynamics of stress-induced alternations of intrinsic amygdala connectivity and neuroendocrine levels. PloS one 10, e0124141 (2015).

30 Vaisvaser, S. et al. Neural traces of stress: cortisol related sustained enhancement of amygdala-hippocampal functional connectivity. Frontiers in human neuroscience 7, 313 (2013).

31 Golden, S. A., Covington III, H. E., Berton, O. & Russo, S. J. A standardized protocol for repeated social defeat stress in mice. Nature protocols 6, 1183–1191 (2011).

32 Krishnan, V. et al. Molecular adaptations underlying susceptibility and resistance to social defeat in brain reward regions. Cell 131, 391–404 (2007).

33 Hodes, G. E. et al. Individual differences in the peripheral immune system promote resilience versus susceptibility to social stress. Proceedings of the National Academy of Sciences 111, 16136–16141 (2014).

34 Hultman, R. et al. Brain-wide electrical spatiotemporal dynamics encode depression vulnerability. Cell 173, 166–180. e114 (2018).

35 Russo, S. J., Murrough, J. W., Han, M.-H., Charney, D. S. & Nestler, E. J. Neurobiology of resilience. Nature neuroscience 15, 1475–1484 (2012).

36 Van Oort, J. et al. How the brain connects in response to acute stress: A review at the human brain systems level. Neuroscience & Biobehavioral Reviews 83, 281–297 (2017).

37 Steiger, J. H. Tests for comparing elements of a correlation matrix. Psychological bulletin 87, 245 (1980).

38 Watanabe, N. & Takeda, M. Neurophysiological dynamics for psychological resilience: A view from the temporal axis. Neuroscience Research 175, 53–61 (2022).

39 Kumar, S. et al. Prefrontal cortex reactivity underlies trait vulnerability to chronic social defeat stress. Nature communications 5, 4537 (2014).

40 Tai, A. P., Leung, M.-K., Geng, X. & Lau, W. K. Conceptualizing psychological resilience through resting-state functional MRI in a mentally healthy population: a systematic review. Frontiers in behavioral neuroscience 17 (2023).

41 Sridharan, D., Levitin, D. J. & Menon, V. A critical role for the right fronto-insular cortex in switching between central-executive and default-mode networks. Proceedings of the National Academy of Sciences 105, 12569–12574 (2008).

42 Capilla, A. et al. The natural frequencies of the resting human brain: an MEG-based atlas. Neuroimage 258, 119373 (2022).

43 Laufs, H. et al. Electroencephalographic signatures of attentional and cognitive default modes in spontaneous brain activity fluctuations at rest. Proceedings of the National Academy of Sciences 100, 11053–11058 (2003).

## Methods references

44 Kong, F., Ma, X., You, X. & Xiang, Y. The resilient brain: psychological resilience mediates the effect of amplitude of low-frequency fluctuations in orbitofrontal cortex on subjective well-being in young healthy adults. Social cognitive and affective neuroscience 13, 755–763 (2018).

45 Miyagi, T. et al. Psychological resilience is correlated with dynamic changes in functional connectivity within the default mode network during a cognitive task. Scientific reports 10, 17760 (2020).

46 Santarnecchi, E. et al. Brain functional connectivity correlates of coping styles. Cognitive, Affective, & Behavioral Neuroscience 18, 495–508 (2018).

47 Windle, G., Bennett, K. M. & Noyes, J. A methodological review of resilience measurement scales. Health and quality of life outcomes 9, 1–18 (2011).

48 Salisu, I. & Hashim, N. A critical review of scales used in resilience research. IOSR Journal of Business and Management 19, 23–33 (2017).

49 Crum, A. J., Salovey, P. & Achor, S. Rethinking stress: the role of mindsets in determining the stress response. J Pers Soc Psychol 104, 716–733 (2013).

50 Cohen, S., Kamarck, T. & Mermelstein, R. A global measure of perceived stress. Journal of health and social behavior 385–396 (1983).

51 Beck, A. T., Steer, R. A. & Carbin, M. G. Psychometric properties of the Beck Depression Inventory: Twenty-five years of evaluation. Clinical psychology review 8, 77–100 (1988).

52 Lewis, A. H., Porcelli, A. J. & Delgado, M. R. The effects of acute stress exposure on striatal activity during Pavlovian conditioning with monetary gains and losses. Frontiers in behavioral neuroscience 8, 179 (2014).

53 Bhanji, J. P., Kim, E. S. & Delgado, M. R. Perceived control alters the effect of acute stress on persistence. Journal of Experimental Psychology: General 145, 356 (2016).

54 Speer, M. E. & Delgado, M. R. Reminiscing about positive memories buffers acute stress responses. Nature human behaviour 1, 0093 (2017).

55 Glover, G. H., Li, T. Q. & Ress, D. Image-based method for retrospective correction of physiological motion effects in fMRI: RETROICOR. Magnetic Resonance in Medicine: An Official Journal of the International Society for Magnetic Resonance in Medicine 44, 162–167 (2000).

56 Kasper, L. et al. The PhysIO toolbox for modeling physiological noise in fMRI data. Journal of neuroscience methods 276, 56–72 (2017).

57 Glasser, M. F. et al. A multi-modal parcellation of human cerebral cortex. Nature 536, 171–178 (2016).

58 Tian, Y., Margulies, D. S., Breakspear, M. & Zalesky, A. Topographic organization of the human subcortex unveiled with functional connectivity gradients. Nature neuroscience 23, 1421–1432 (2020).

59 Rolls, E. T., Huang, C.-C., Lin, C.-P., Feng, J. & Joliot, M. Automated anatomical labelling atlas 3. Neuroimage 206, 116189 (2020).

60 Sorensen, T. A method of establishing groups of equal amplitude in plant sociology based on similarity of species content and its application to analyses of the vegetation on Danish commons. Biologiske skrifter 5, 1–34 (1948).

61 Jafri, M. J., Pearlson, G. D., Stevens, M. & Calhoun, V. D. A method for functional network connectivity among spatially independent resting-state components in schizophrenia. Neuroimage 39, 1666–1681 (2008).

62 Benjamini, Y. & Hochberg, Y. Controlling the false discovery rate: a practical and powerful approach to multiple testing. Journal of the Royal statistical society: series B (Methodological) 57, 289–300 (1995).

63 Yarkoni, T., Poldrack, R. A., Nichols, T. E., Van Essen, D. C. & Wager, T. D. Large-scale automated synthesis of human functional neuroimaging data. Nature methods 8, 665–670 (2011).

64 Allen, P. J., Josephs, O. & Turner, R. A method for removing imaging artifact from continuous EEG recorded during functional MRI. Neuroimage 12, 230–239 (2000).

65 Sedghamiz, H. Matlab implementation of Pan Tompkins ECG QRS detector. Code Available at the File Exchange Site of MathWorks (2014).

66 Pan, J. & Tompkins, W. J. A real-time QRS detection algorithm. IEEE transactions on biomedical engineering, 230–236 (1985).

