## SupplementaryInformation for "Neural signatures of human psychological resilience driven by acute stress"

**Supplementary figures and legends**

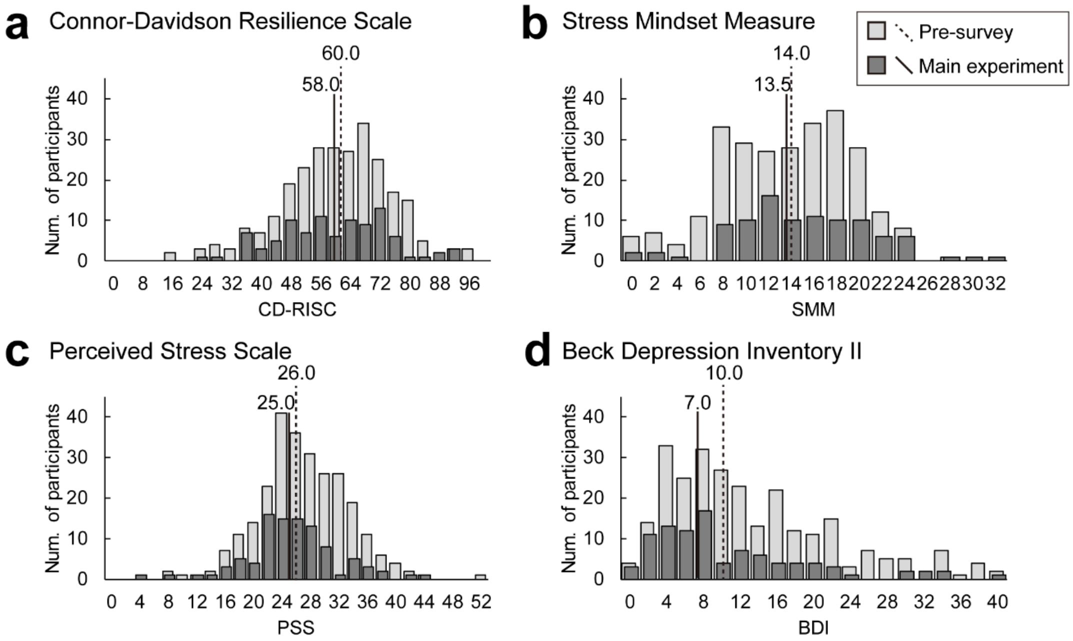


**Supplementary Fig. 1: Score distribution of stress-related questionnaires.**

Prior to collecting neurophysiological data, a pre-survey was conducted to understand the distribution of (**a**) the Connor-Davidson Resilience Scale (CD-RISC)[4], (**b**) the Stress Mindset Measure (SMM)[41], (**c**) the Perceived Stress Scale (PSS)[42], and (**d**) the Beck Depression Inventory-II (BDI-II)[43]. In total, 265 samples (166 males; mean age, 18–24 years) were collected from undergraduate and graduate students at a Japanese university. The median and inter-quantile range was 60.0 (50.0~69.0) for CD-RISC, 14.0 (9.0~17.0) for SMM, 26.0 (23.0–31.0) for PSS, and 10.0 (6.0–17.0) for BDI. Tests of normality for each trait showed that the distribution of the CD-RISC did not deviate from the normal distribution ($D$ = 0.052, $P_{FDR}$ = 0.085), but the other traits were not normally distributed (SMM: $D$ = 0.091, $P_{FDR}$ = 3.017E-5; PSS: $D$ = 0.057, $P_{FDR}$ = 0.047; BDI: $D$ = 0.123, $P_{FDR}$ = 7.738E-10). In the neurophysiological experiment (N = 96, 72 males), the median and inter-quantile ranges were 58.0 (48.0~69.0) for CD-RISC, 13.5 (10.8~19.0) for SMM, 25.0 (21.8~28.0) for PSS, and 7.0 (4.0~14.0) for BDI. The two-sample Kolmogorov–Smirnov test showed that there was no sampling bias between the pre-survey and the main experimental population in any of the questionnaires (CD-RISC: $Z$ = 0.794, $P_{FDR}$ = 0.654; SMM: $Z$ = 0.889, $P_{FDR}$ = 0.545; PSS: $Z$ = 1.230, $P_{FDR}$ = 0.194; BDI: Z = 1.490, $P_{FDR}$ = 0.095). Therefore, we were successful in randomly sampling the general population for the neurophysiological experiments.

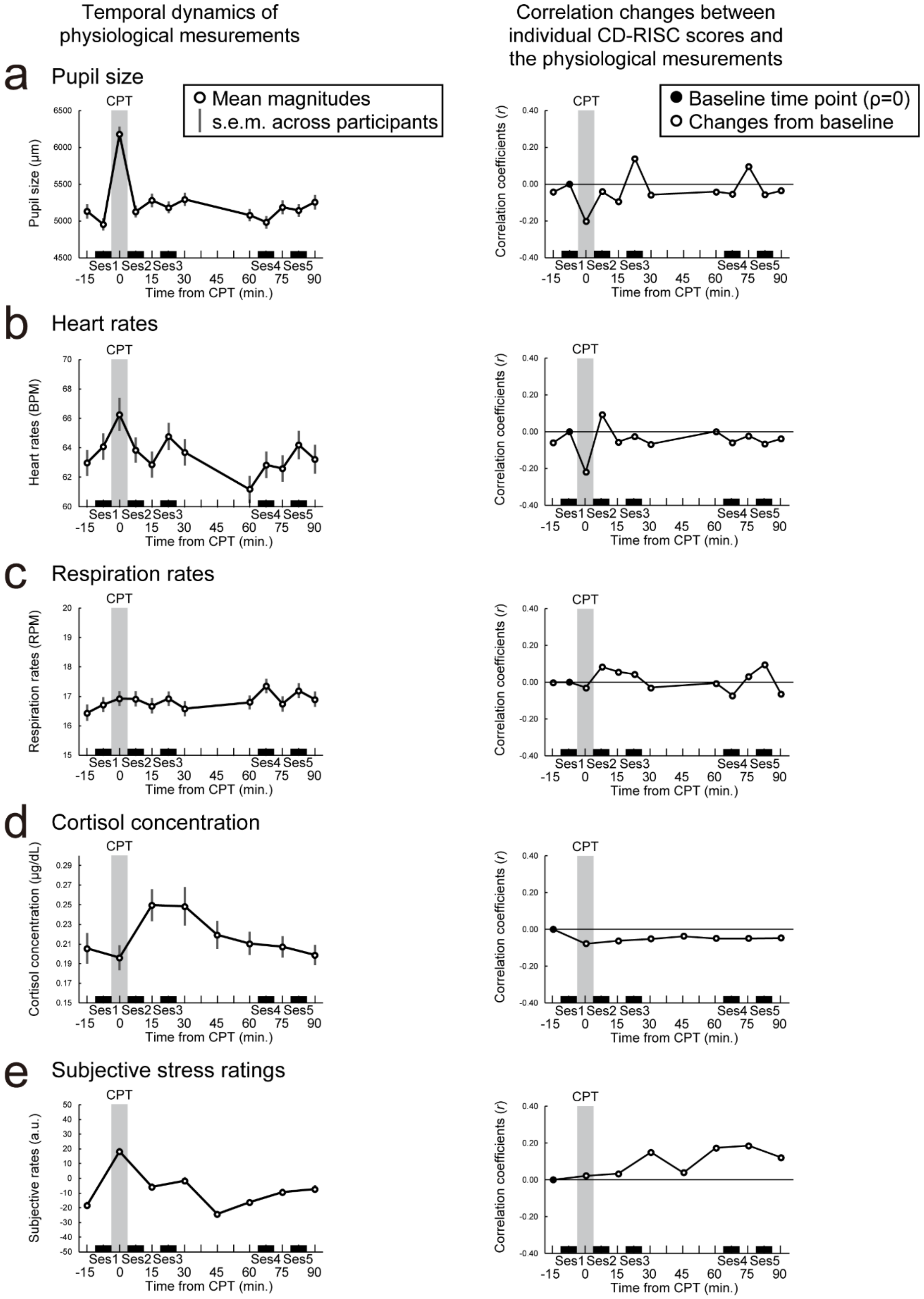


Supplementary Fig. 2: Temporal dynamics of peripheral and subjective responses to acute stress and their correlations with CD-RISC.

Pupil size (μm), heart rate (beats per minute: BPM), respiration rate (respirations per minute: RPM), cortisol concentration density (μg/dL), and subjective stress ratings (rating from -50 to 50) were collected several times during the experiments. The left column shows the temporal dynamics of each measurement; right column, temporal correlations between the CD-RISC and each measurement. Correlations with the CD-RISC were performed after each physiological value was calculated as the change from the pre-stressed baseline (shown as a black dot). We observed significant changes in pupil size, heart rate, cortisol levels, and subjective ratings following acute stress exposure. However, individual differences in these responses did not correlate with individual levels of resilience. (**a**) Pupil size at CPT (shown as shaded grey area) was significantly increased compared to the other time points (Wilcoxon signed-rank sum test: $z$ = 7.368~8.193, $P_{FDR}$ = 8.71E-15~81.04E-12, $r$ = 0.813~0.869), but these changes in pupil size from baseline (shown as the black dot) showed no significant correlation with individual CD-RISC scores (Spearman correlation $\rho$ = -0.203~0.139, $P_{FDR}$ = 0.741~0.750). (**b**) Similarly, heart rates during CPT were significantly increased compared to those at other time points (Wilcoxon signed-rank sum test: $z$ = 2.642–6.463, $P_{FDR}$ = 1.36E-9–0.012, $r$ = 0.273–0.663), except for Session 3 (CPT vs. Session 3: $z$ = 1.592, $P_{FDR}$ = 0.136, $r$ = 0.164). However, correlations between CD-RISC and changes in heart rate did not show significant correlations at any time points (Spearman correlation: $\rho$ = -0.219~0.094, $P_{FDR}$ = 0.399~0.992). (**c**) Mean respiration rates at CPT were not changed compared to the other time points (Wilcoxon signed-rank sum test: $z$ = 0.004~2.189, $P_{FDR}$ = 0.100~0.997, $r$ = 0.000~0.223), nor in the correlations with CD-RISC (Spearman correlation: $\rho$ = -0.072~0.097, $P_{FDR}$ = 0.955~0.999). (**d**) Cortisol densities increased from 15 to 30 min after stress exposure, and returned to baseline densities by 60 min (Wilcoxon signed-rank sum test at 15 min: $z$ = 1.995~4.859, $P_{FDR}$ = 1.15E-6~0.046, $r$ = 0.233~0.561; at 60 min: $z$ = 1.885~3.348, $P_{FDR}$ = 0.001~0.059, $r$ = 0.216~0.389), but there were no significant correlations with CD-RISC (Spearman correlation: $\rho$ = -0.077~ -0.036 $P_{FDR}$ = 0.742). (**e**) Subjective stress ratings confirmed that stress levels peaked during the CPT (Wilcoxon signed-rank sum test: $z$ = 5.853~8.312, $P_{FDR}$ = 2.80E-15~9.65E-9, $r$ = 0.601~0.853), but did not correlate with CD-RISC (Spearman correlation: $\rho$ = -0.009~ -0.208 $P_{FDR}$ = 0.288~0.993).

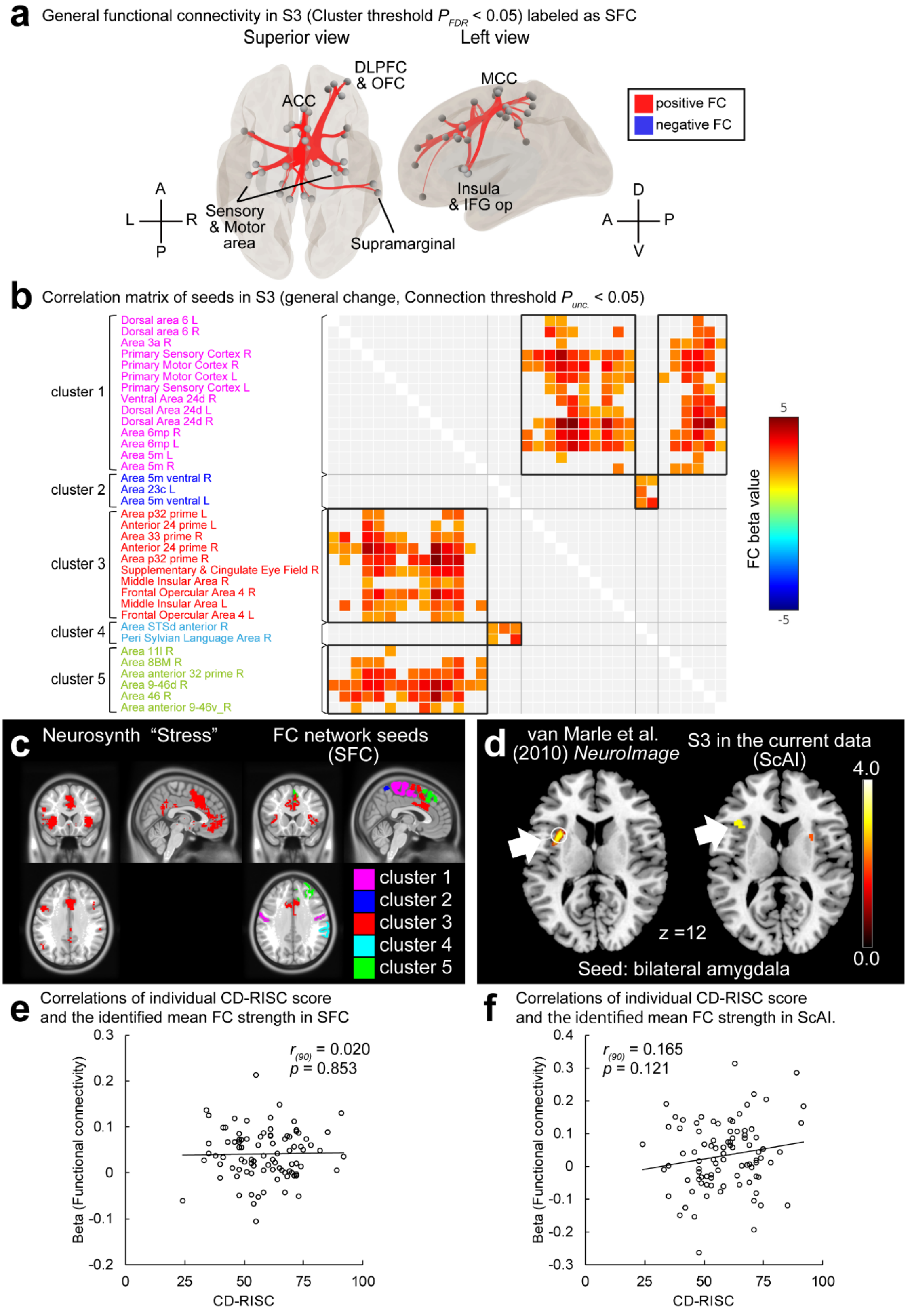


**Supplementary Fig. 3: Stress-driven functional connectivity network identified in**

**S3.**

Changes in functional connectivity (FC) after acute stress from the baseline session (labelled S2 to S5) were compared with those in previous fMRI studies. These validation analyses confirmed that our current experiment successfully induced stress-related brain activation in S3 and allowed us to assess the relationships between stress-related neural responses and individual resilience levels. (**a**) Stress-related functional connectivity analysis showing increased activation in the anterior and medial cingulate, bilateral sensorimotor, and bilateral insular cortices approximately 20 min after stress exposure (S3), labelled SFC. Cluster threshold $P_{FDR} < 0.05$. (**b**) The coloured grids in the matrix indicate significant correlations between two seeds (cluster threshold $P_{unc.} < 0.05$). Seed names are listed according to the classification of Glasser (2016)[57]. (**c**) The left panel shows the result of the brain activation map based on the meta-analysis of 321 studies related to ‘stress’ (uniformity test $P < 0.01$, FDR-corrected) using Neurosynth[63]. The right panel shows a seed map of the SFC as shown in (**a**). The functional seed regions in the current study overlapped with those in the previous meta-analysis. (**d**) Stress-related amygdala-insula connection (ScAI). We performed the same analysis as van Marle et al., who reported that the FC between the bilateral amygdala and the left anterior insular cortex increased 20 min after stress exposure[20]. Consistent with this report, we identified a cluster from the left anterior insular cortex to the inferior frontal operculum at S3. Peak coordinates [x, y, z] = -40, 22, 12: $T_{(90)}$ = 4.18, Cluster size k = 30, $P_{unc.}$ = 3.35E-05. (**e, f**) Scatter plots of individual CD-RISC scores and identified SFC (**e**) or ScAI (**d**). No significant correlations were observed between the CD-RISC and stress-related brain activation. The image on the left in (**d**) was reproduced from van Marle et al.[20] with permission from Elsevier.

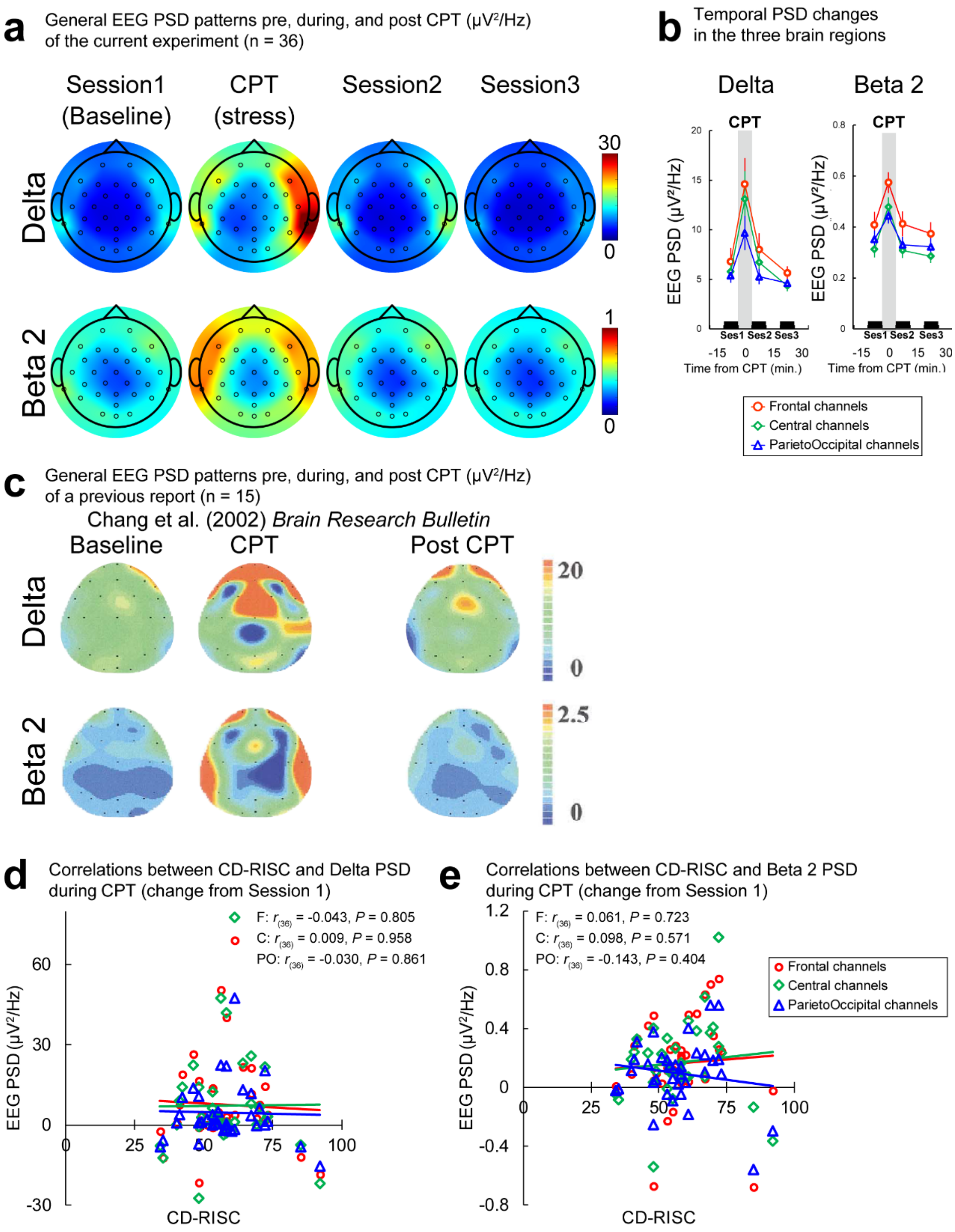


**Supplementary Fig. 4: Spectral analysis of EEG driven by acute stress.**

The temporal dynamics of the EEG power spectral density (PSD) showed that the power in the delta frequency band (0.5–3.5 Hz) and the high-beta frequency band (beta

2: 21.5–29.5 Hz) increased during the cold pressor test (CPT). Note that the topographic maps presented here show the PSD at each session in line with the description of previous studies, and not the difference from the pre-stressed baseline. (**a**) Temporal changes in topographic maps from Session 1, CPT, Session 2, and Session 3 for delta and beta 2 components. (**b**) In the delta band, the mean PSDs increased during CPT, especially in the frontal and parieto-occipital channels (paired t-test of Session 1 vs. CPT in the frontal channels: $T_{(35)}$ = -2.650, $P_{FDR}$ = 0.036, $r$ = 1.409), and decreased in Session 3 (paired t-test of CPT vs. Session 3 in the frontal channels: $T_{(35)}$ = 3.239, $P_{FDR}$ = 0.016, $r$ = 1.480; in parieto-occipital channels: $T_{(35)}$ = 2.909, $P_{FDR}$ = 0.038, $r$ = 1.441). Similarly, in the delta band, the mean PSDs in the frontal, central, and parieto-occipital channels increased at CPT (Paired t-test of Session 1 vs. CPT in the frontal channels: $T_{(35)}$ = -3.259, $P_{FDR}$ = 0.005, $r$ = 0.483; in the central channels: $T_{(35)}$ = -3.788, $P_{FDR}$ = 0.001 $r$ = 0.539; in the parieto-occipital channels: $T_{(35)}$ = -2.557, $P_{FDR}$ = 0.030, $r$ = 0.397) and decreased in Session 2 (Paired t-test of CPT vs. Session 2 in the frontal channels: $T_{(35)}$ = 3.464, $P_{FDR}$ = 0.004, $r$ = 0.505; in Central channels: $T_{(35)}$ = 3.819, $P_{FDR}$ = 0.001 $r$ = 0.542; in the parieto-occipital channels: $T_{(35)}$ = 3.373, $P_{FDR}$ = 0.007, $r$ = 0.495). (**c**) The topographic map shown was obtained from Chang et al.[13]. The observed PSD changes in delta and beta 2 during CPT, shown in (**a**), are consistent with these reports and those of other groups[14,15]. (**d, e**) The scatter plots of individual CD-RISC scores and identified stress-related PSD delta (**d**) and beta 2 (**e**). No significant correlations were observed between the CD-RISC and the identified stress-related PSD in any channel. (**c**) Reproduced from Chang et al.[12] with permission from Elsevier.

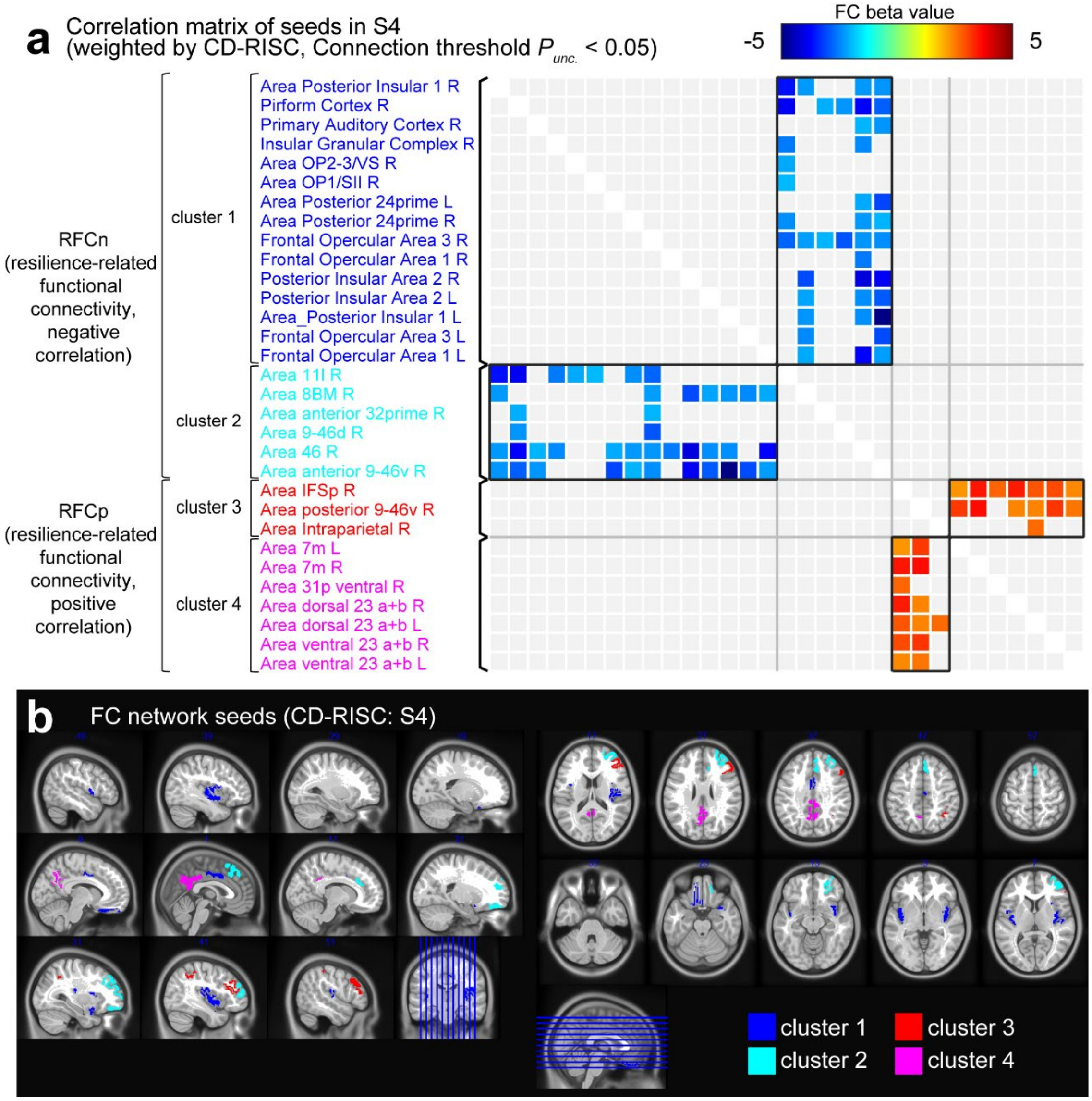


**Supplementary Fig. 5: Seeds of the resilience-related functional connectivity network, labelled RFCn and RFCp.**

(**a**) Correlation matrix of seeds in S4. The coloured grids in the matrix indicate significant correlations between the two seeds (cluster threshold, $P_{unc.} < 0.05$). Seed names are listed according to the classification of Glasser (2016)[57]. (**b**) Seed maps of RFCn (clusters 1 and 2) and RFCp (clusters 3 and 4).

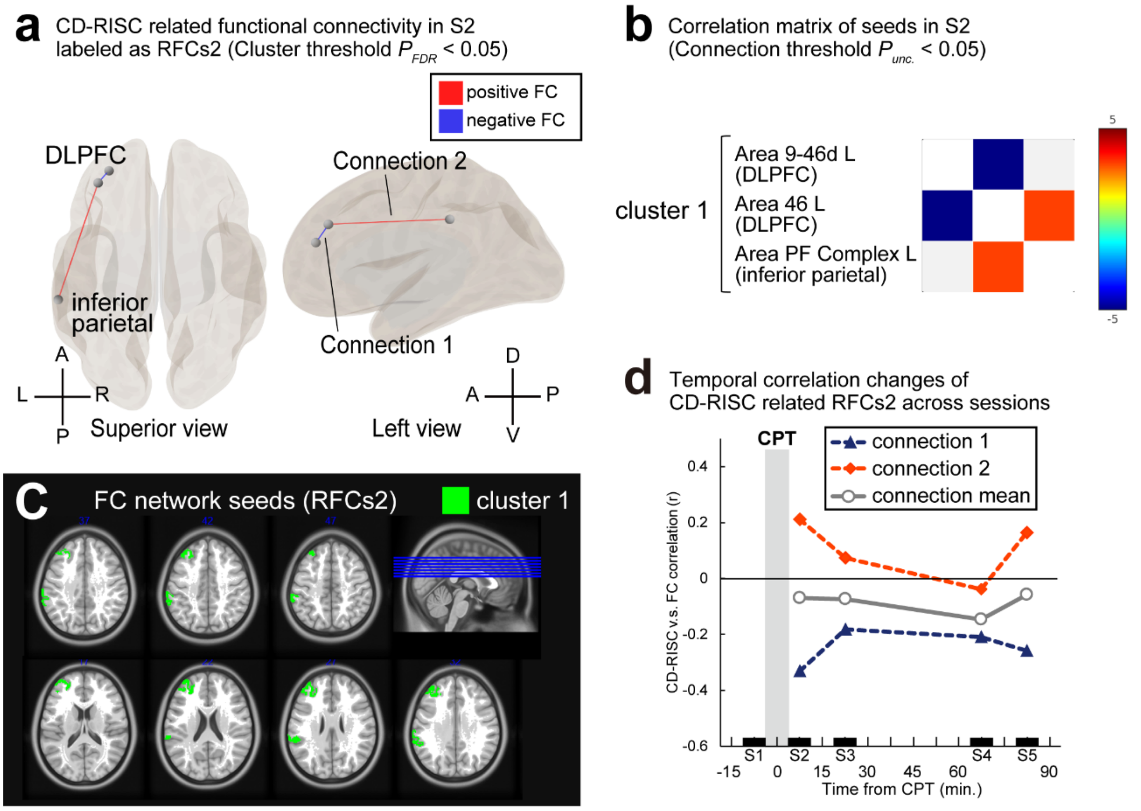


**Supplementary Fig. 6: fMRI functional connectivity correlated with resilience immediately after acute stress experience.**

(**a**) A significant FC was identified in S2 with a cluster-level threshold $P_{FDR}$ of < 0.05. This cluster included both positive and negative connections with CD-RISC scores (RFCs2). Seeds were distributed in the left dorsolateral prefrontal and inferior parietal cortices. (**b**) Correlation matrix of seeds in S2. The coloured grids in the matrix indicate significant correlations between two seeds (cluster threshold, $P_{unc.}$ < 0.05), and (**c**) shows the seed map of RFCs2. (**d**) Temporal dynamics of the correlation between CD-RISC and RFCs2. Since RFCs2 includes both positive and negative connections, the graph shows the change in the correlation coefficients between the CD-RISC and individual connections, in addition to the cluster. The correlation coefficient (*r*) of the connections did not significantly change between S2 and S5 ($P_{FDR} < 0.05$, corrected).

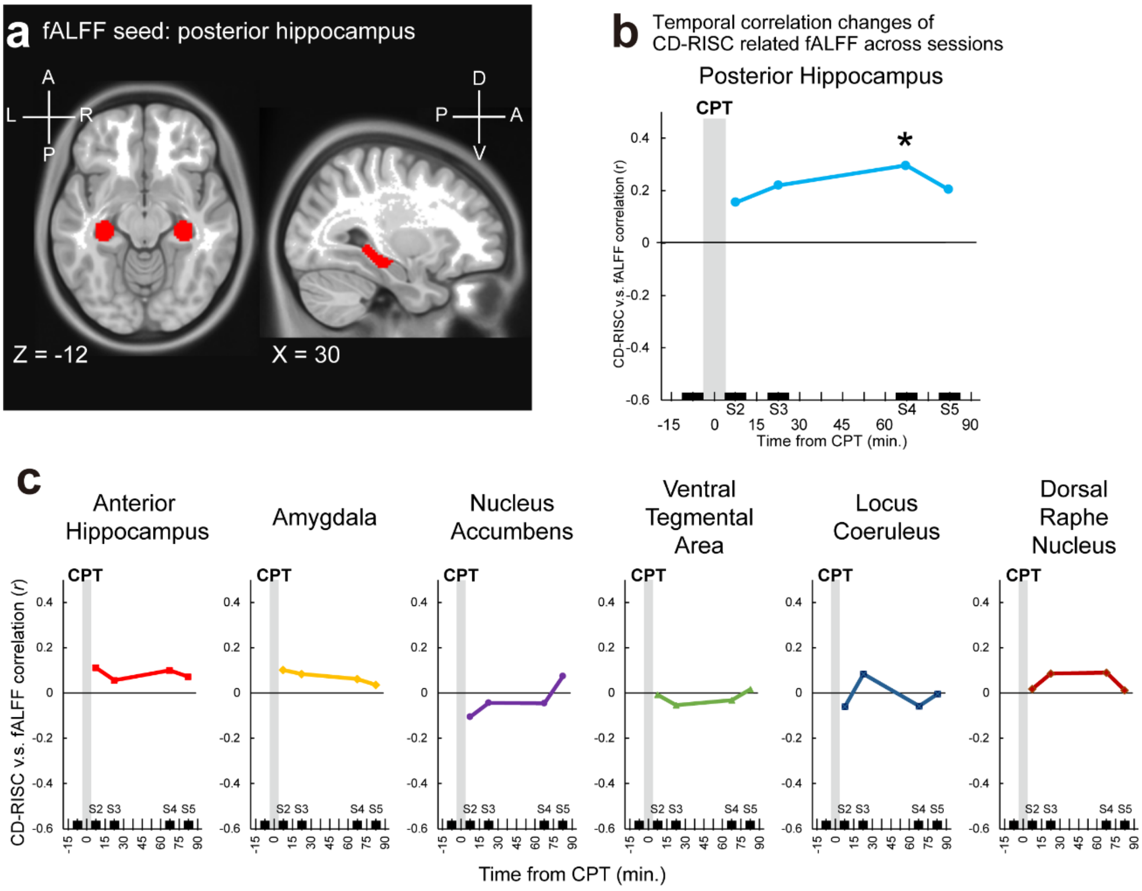


**Supplementary Fig. 7: Temporal dynamics of fractional amplitude of low-frequency fluctuations in the selected subcortical regions.**

Spontaneous neural activity correlating with individual CD-RISC scores was evaluated using the fractional amplitude of low-frequency fluctuations (fALFF). Seven regions of interest (ROIs), the bilateral anterior hippocampus, posterior hippocampus, amygdala, nucleus accumbens, ventral tegmental area, locus coeruleus, and dorsal raphe nucleus, were selected based on a previous review of resilience[38]. We found a significant correlation between individual CD-RISC scores and fractional amplitudes only in the posterior hippocampus (HIPp) at S4. (**a**) HIPp seed map (**b**) Temporal dynamics of the correlation between CD-RISC and fALFF in HIPp. The positive correlation with the CD-RISC scores peaked in S4 ($r_{(90)} = 0.296$, $P_{FDR} = 0.033$), but the correlation coefficients did not show significant differences from S2 to S5 (Hotelling–Williams test: $T_{(90)} = -1.391\sim0.972$, $P_{FDR} = 0.787\sim0.809$). (**c**) Temporal dynamics of the CD-RISC scores and obtained ROIs. No significant correlations were observed in the other ROIs ($r_{(90)} = -0.106\sim0.112$, $P_{FDR} = 0.591\sim0.973$).

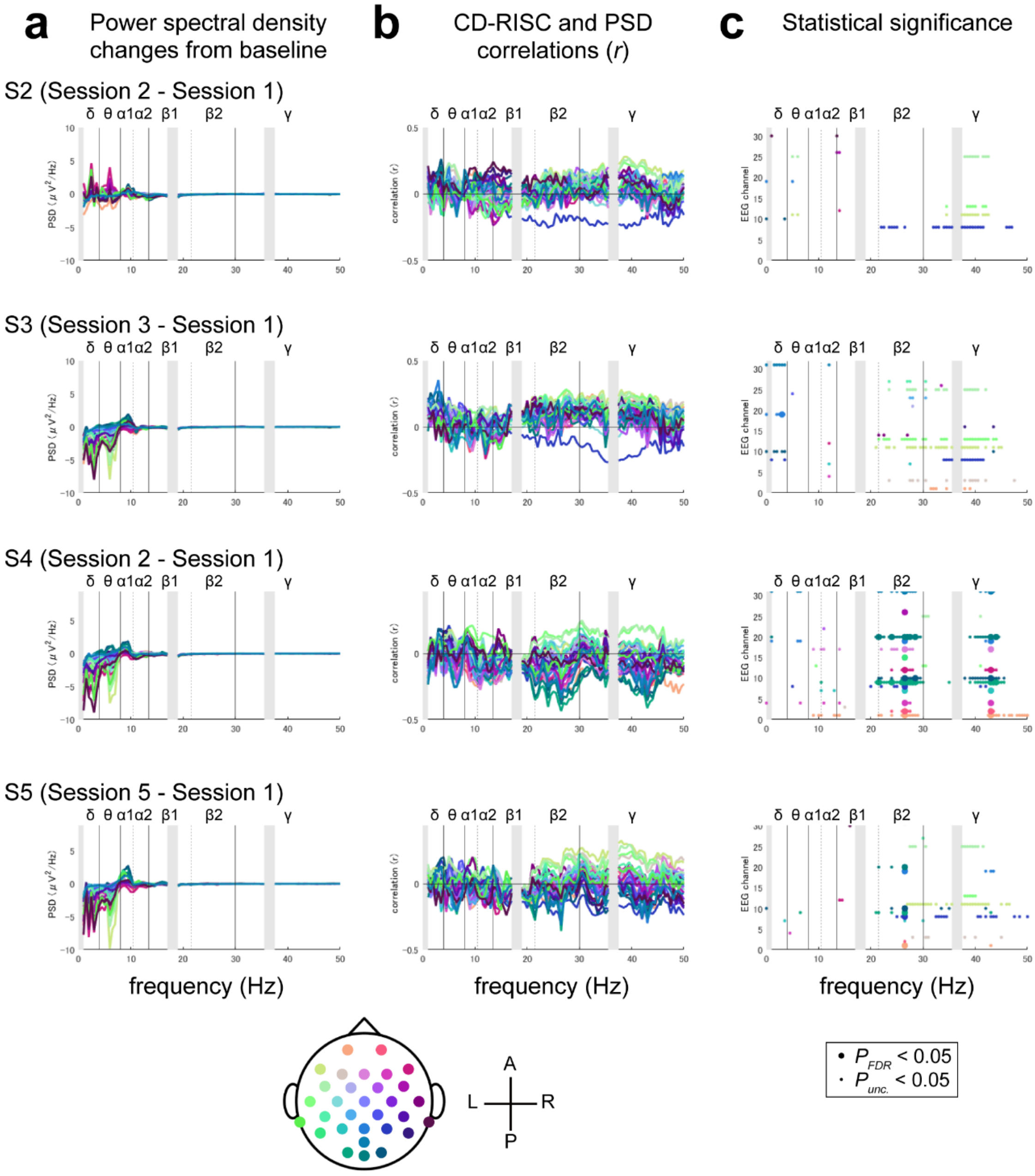


945 **Supplementary Fig. 8: Spectral analyses of EEG from S2 to S5 and their**

946 **correlations with individual CD-RISC scores.**

947 (**a**) Changes in power spectral density (PSD) in each session from baseline Session one

948 to 0.5 Hz. (**b**) Correlation coefficients between individual CD-RISC scores and PSD for

949 each frequency band. (**c**) Statistically significant frequencies for each channel. Large

dots in the graph indicate a significant channel and frequency at the threshold with $P_{FDR} < 0.05$, whereas small dots indicate the threshold with $P_{unc.} < 0.05$. The colour of each line corresponds to the 31 channels located on the scalp. Grey masks represent bands that were not detectable owing to noise from the fMRI scanner.

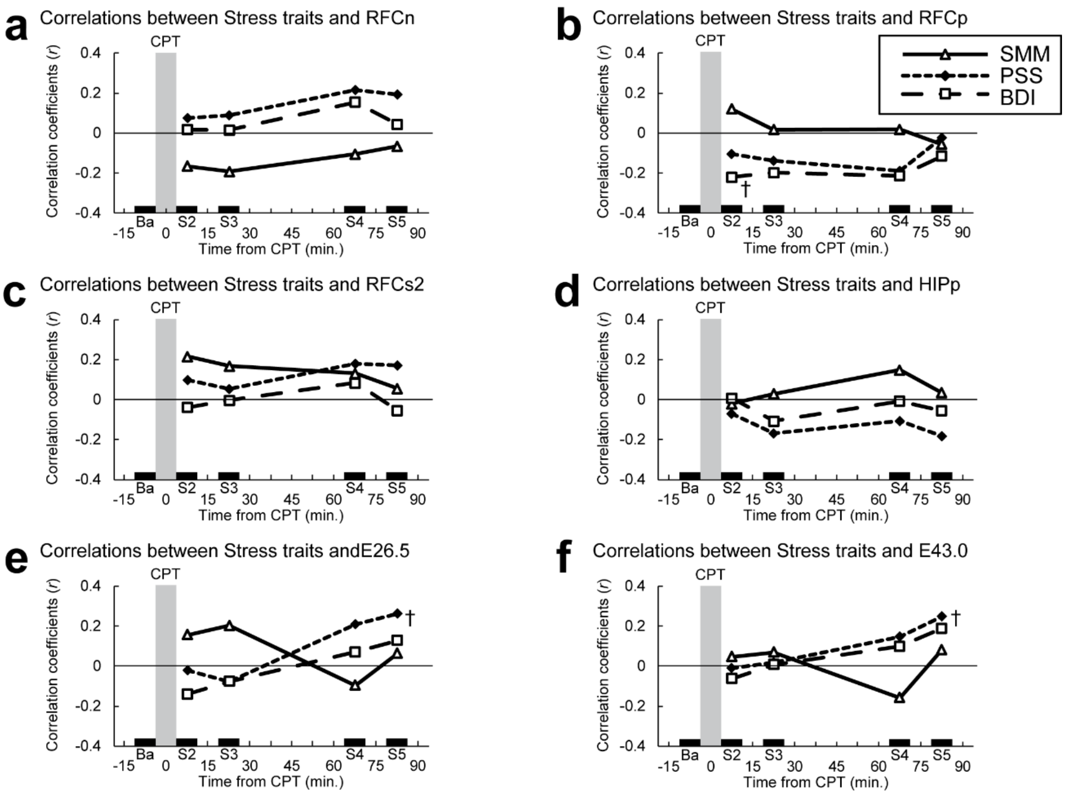


**Supplementary Fig. 9: Temporal dynamics of the correlation between the three stress trait scores and resilience-related fMRI or EEG activities**

These graphs show the correlation between the three stress-related personality traits and the temporal dynamics of the six brain activities identified in the current study. In summary, none of the brain activities significantly correlated with individual scores on the SMM, PSS, and BDI. $^{\dagger}P_{FDR} < 0.10$. (**a**) The ranges of correlation coefficients and P-values with RFCn were $r_{(90)}$ = -0.192~0.217 and $P_{FDR}$ = 0.135~0.881. (**b**) The ranges of correlation coefficients and P-values with RFCp were $r_{(90)}$ = -0.219~0.121 and $P_{FDR}$ = 0.083~0.869. (**c**) The ranges of correlation coefficients and P-values with RFCs1 were $r_{(90)}$ = -0.053~0.216 and $P_{FDR}$ = 0.165~0.982. (**d**) The range of correlation coefficients and P-values with HIPp were $r_{(90)}$ = -0.181~0.149 and $P_{FDR}$ = 0.233~0.944. (**e**) The ranges of correlation coefficients and P-values with RFCn were E26.5($r_{(88)}$ = -0.140~0.261 and $P_{FDR}$ = 0.052~0.844. (**f**) The ranges of correlation coefficients and P-values with RFCn were E43.0($r_{(88)}$ = -0.157~0.249 and $P_{FDR}$ = 0.072~0.940.

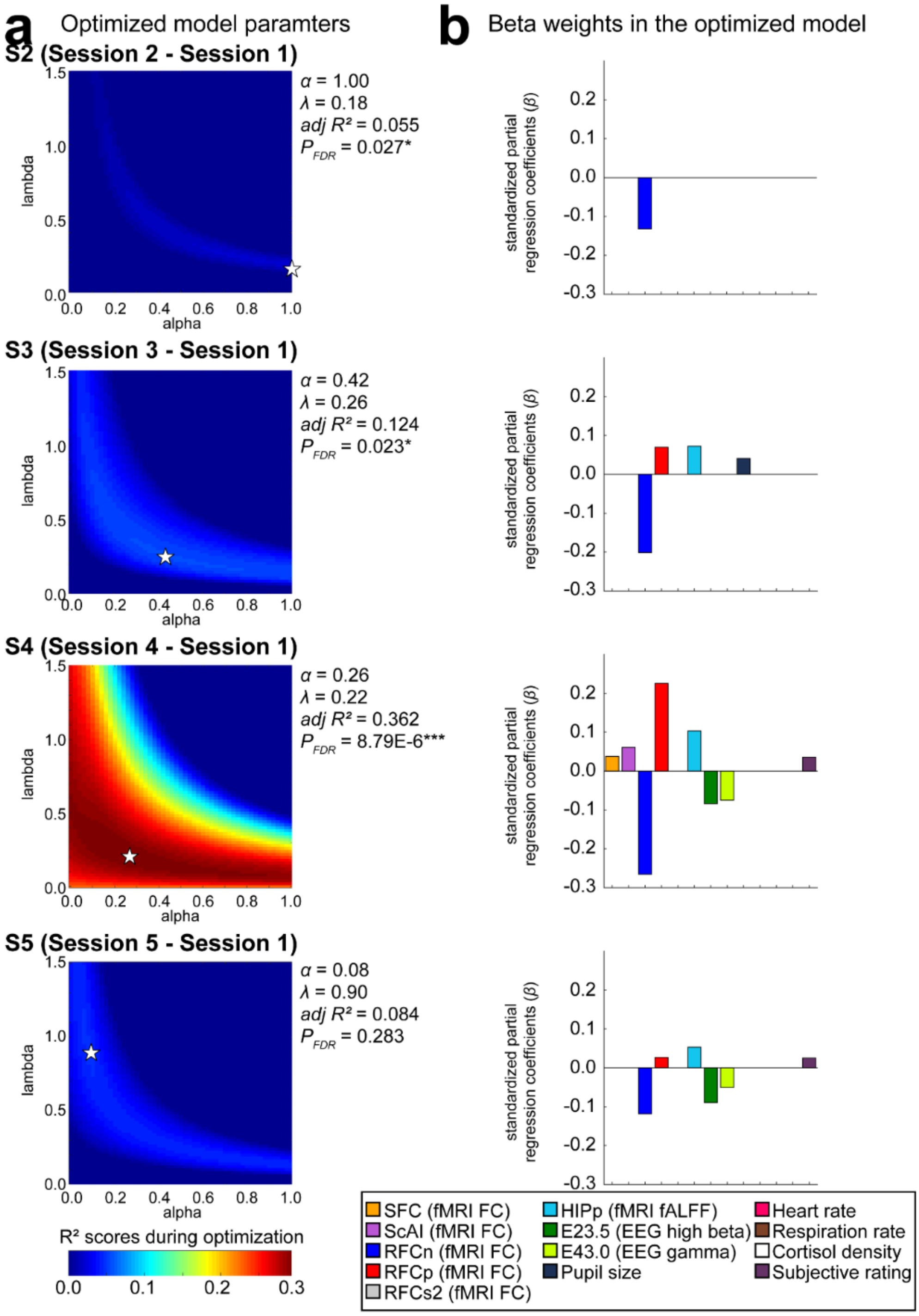


**Supplementary Fig. 10: The optimised combination of weights to predict individual resilience levels in each session.**

Model 2 was optimised using 13 factors in each session. (**a**) Heat map showing the $R^2$

values during optimisation of the alpha and lambda parameters using an elastic net. The stars on the map indicate the peak values. (**b**) Estimated weights of selected factors to predict individual CD-RISC scores. The optimised model in S4 had the largest $R^2$ value across the four sessions. Consistent with Model 1, as shown in Fig. 4, RFCn and RFCp contributed dominantly to the prediction of individual resilience levels in S4.

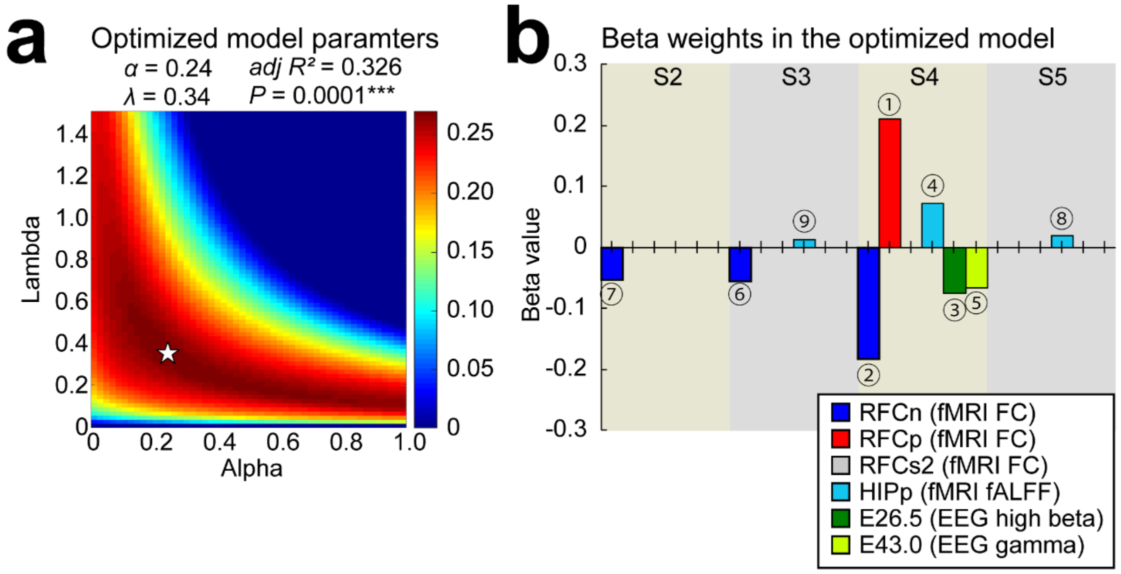


**Supplementary Fig. 11: The optimised combination of weights to predict individual resilience levels using the six resilience-related factors.**

Model 3 was optimised using the six neural factors identified in this study. (**a**) Heat map showing the $R^2$ values during optimisation of the alpha and lambda parameters using an elastic net. The stars on the map indicate the peak values. (**b**) Estimated weights of selected factors to predict individual CD-RISC scores. Consistent with Model 1, as shown in Fig. 4, RFCn and RFCp in S4 dominantly contributed to the prediction of individual resilience levels.

**Supplementary Discussion 1: Validation of stress-induced peripheral and neural responses**

Before exploring resilience-related responses, we confirmed the efficacy of acute stress manipulation through peripheral and brain responses. Consistent with previous reports[6-9,16-19], CPT successfully induced peripheral autonomic nervous system responses during stress induction, except for changes in respiration (Supplementary Fig. 2a-c left)[11]. Salivary cortisol levels also increased at 15 and 30 min post-stress. Subjective measures of stress peaked during the CPT (Supplementary Fig. 2d,e, left).

CPT also successfully induced brain responses that were spatially and temporally consistent with those reported previously. First, we observed changes in functional connectivity (FC) using fMRI before (baseline) and after acute stress exposure (labelled S2–S5). The stress-driven functional connectivity (SFC) network in the anterior and medial cingulate, sensory-motor, and insular cortices increased in S3 (Supplementary Fig. 3a,b; cluster-level $P_{FDR} < 0.05$). The network of regions identified in S3 overlapped with the network of stress-related regions observed in the Neurosynth meta-analysis[63] using the term ‘stress’ (Supplementary Fig. 3c). In addition, a previous resting-state fMRI study reported that stress increased the FC between the bilateral amygdala and the left anterior insular cortex[20]. Therefore, we conducted the same analysis seeded in the bilateral amygdalae and identified a similar cluster in the left anterior insular cortex (but partially extended to the inferior frontal operculum) in S3 (Supplementary Fig. 3d, named ScAI; the stress-related connection between the amygdala and insula).

Second, with respect to the EEG, we were able to replicate previous results showing that EEG PSD in the delta frequency band (0.5–3.5 Hz) and a part of the beta frequency band (21.5–29.5 Hz; beta 2) increased during CPT[13-15] (Supplementary Fig. 4a-c, $P_{FDR} < 0.05$). Therefore, like the peripheral responses, the fMRI and EEG results indicated that our experimental design successfully induced neural activation driven by an acute stressor.

**Supplementary Discussion 2: Functional roles of identified factors**

The RFCn and RFCp networks identified in S4 strongly overlapped with the salience network (SaN) and default mode network (DMN), respectively[36]. Hermans et al.[28] argued that SaN increases within an hour of acute stress and decreases as a dynamic equilibrium after an hour. The identified RFCn suggests that the decrease in SaN after 1 hour was more pronounced in the resilient individuals. In contrast, the DMN (RFCp) was increased in resilient individuals. Although increased numbers of DMN after acute stress exposure have been repeatedly reported, their functional role has been debated for 2 decades[36]. Our results suggest that a gradual increase in the DMN after stress exposure contributes to the expression of psychological resilience.

Similar to increased activation in the DMN, spontaneous hippocampal activation was positively correlated with resilience scores. The hippocampus is also considered part of the DMN[36], and this activity could be attributed to the DMN. Hippocampal activity was observed in the posterior region of the brain. The anterior part of the hippocampus is more important for the stress response than the posterior part, and the posterior part is more involved in memory consolidation[67]. Considering that appropriate suppression and consolidation of past stressful memories are important for the formation of resilience[68], individual sensitivity to memory consolidation may be affected by posterior hippocampal activation[69].

Furthermore, frequency analysis of the EEG revealed that the power of the high-beta and gamma components increased over 60 min after stress in less resilient individuals. High-beta power increases during stress exposure[13-15]. We showed that high-beta power increased 1 hour after stress in less resilient individuals. Clinical studies using magnetoencephalography (MEG) have reported similar increases in beta power in patients with depression and PTSD patients[70,71]. Compared to the findings on beta bands, the relationship between gamma bands and resilience remains unclear. While some reports are consistent with our findings and show an increase in prefrontal-limbic resting gamma power in patients with depression and PTSD[72,73], others have reported a decrease in gamma power in patients with depression[74]. Thus, the role of gamma power in resilience requires further investigation.

### References for Supplementary Discussion


67 Fanselow, M. S. & Dong, H.-W. Are the dorsal and ventral hippocampus functionally distinct structures? *Neuron* 65, 7-19 (2010).

68 Mary, A. et al. Resilience after trauma: The role of memory suppression. *Science* 367, eaay8477 (2020).

69 Schmidt, M. V. et al. Individual stress vulnerability is predicted by short-term memory and AMPA receptor subunit ratio in the hippocampus. *Journal of Neuroscience* 30, 16949-16958 (2010).

70 Brunetti, M. et al. Resilience and cross-network connectivity: a neural model for post-trauma survival. *Progress in Neuro-Psychopharmacology and Biological Psychiatry* 77, 110-119 (2017).

71 Nugent, A. C., Robinson, S. E., Coppola, R., Furey, M. L. & Zarate Jr, C. A. Group differences in MEG-ICA derived resting state networks: application to major depressive disorder. *Neuroimage* 118, 1-12 (2015).

72 Dunkley, B. et al. Resting-state hippocampal connectivity correlates with symptom severity in post-traumatic stress disorder. *NeuroImage: Clinical* 5, 377-384 (2014).

73 Jiang, H., Tian, S., Bi, K., Lu, Q. & Yao, Z. Hyperactive frontolimbic and frontocentral resting-state gamma connectivity in major depressive disorder.

*Journal of Affective Disorders* 257, 74-82 (2019).

74 Pizzagalli, D. A., Peccoralo, L. A., Davidson, R. J. & Cohen, J. D. Resting anterior cingulate activity and abnormal responses to errors in subjects with elevated depressive symptoms: A 128-channel EEG study. *Human brain mapping* 27, 185-201 (2006).
